# Nanomolar, noncovalent antagonism of hedgehog cholesterolysis: exception to the “irreversibility rule” for protein autoprocessing inhibition

**DOI:** 10.1101/2021.09.01.458638

**Authors:** Andrew G Wagner, Robert T Stagnitta, Zihan Xu, John L Pezzullo, Nabin Kandel, José-Luis Giner, Douglas F Covey, Chunyu Wang, Brian P Callahan

**Affiliations:** Department of Chemistry, Binghamton University, State University of New York, Binghamton, New York 13902, United States; Department of Chemistry, State University of New York College of Environmental Science and Forestry, Syracuse, New York 13210, United States; Department of Biological Sciences, Center for Biotechnology and Interdisciplinary Studies, Rensselaer Polytechnic Institute, 110 8th Street, Troy, New York 12180, United States; Department of Developmental Biology, Taylor Family Institute for Innovative Psychiatric Research, 660 South Euclid Avenue, St. Louis, Missouri 63110, United States

## Abstract

Hedgehog (Hh) signaling ligands undergo carboxy terminal sterylation through specialized autoprocessing, called cholesterolysis. Sterylation is brought about intramolecularly in a single turn-over by an enzymatic domain, called HhC. HhC is found in precursor Hh proteins only. Through cholesterolysis, HhC is cleaved from the precursor. Attempts to identify molecules that inhibit intramolecular cleavage/sterylation activity of HhC have resulted in antagonists that bind HhC irreversibly through covalent mechanisms, as is commonplace for protein autoprocessing inhibitors. Here we report an exception to the “irreversibility rule” for protein autoprocessing inhibition. Using a FRET-based activity assay for HhC, we screened a focused library of sterol-like analogs for HhC cholesterolysis inhibitors. We identified and validated four structurally related noncovalent inhibitors, which were then used for SAR studies. The most effective derivative, tBT-HBT, binds HhC reversibly with an IC_50_ of 300 nM. An allosteric binding site for tBT-HBT, encompassing interactions from the two subdomains of HhC, is suggested by kinetic analysis, mutagenesis studies, and photoaffinity labeling. A striking resemblance is found between the inhibitors described here and a family of noncovalent, allosteric *activators* of HhC, which we described previously. The inhibitor/activator duality appears to be mediated by the same allosteric site, which displays sensitivity to subtle differences in the structure of a heterocycle substituent on the effector molecule.

## INTRODUCTION

Hedgehog (Hh) ligands, the extracellular signaling factors involved in embryo development and many types of cancers^*1–6*^ appear unique in their post-translational modification by cholesterol.^*7, 8*^ Lipidation occurs by an autoprocessing reaction called cholesterolysis during Hh protein transit through the endoplasmic reticulum (ER). A partially characterized ~25 kDa domain, HhC, present in precursor forms of Hh serves as the intramolecular catalyst for cholesterolysis (**Figure 1**). HhN-chol, the cholesterylated product, is released from HhC and undergoes additional lipidation as it matures into the Hh ligand.^*9*^ Mutations in HhC that suppress cholesterolysis result in retention of the Hh precursor in the ER.^*10, 11*^ Retained precursor is degraded by the ER proteostasis system thereby suppressing downstream Hh signaling.^*10, 12, 13*^

**Figure 1.**
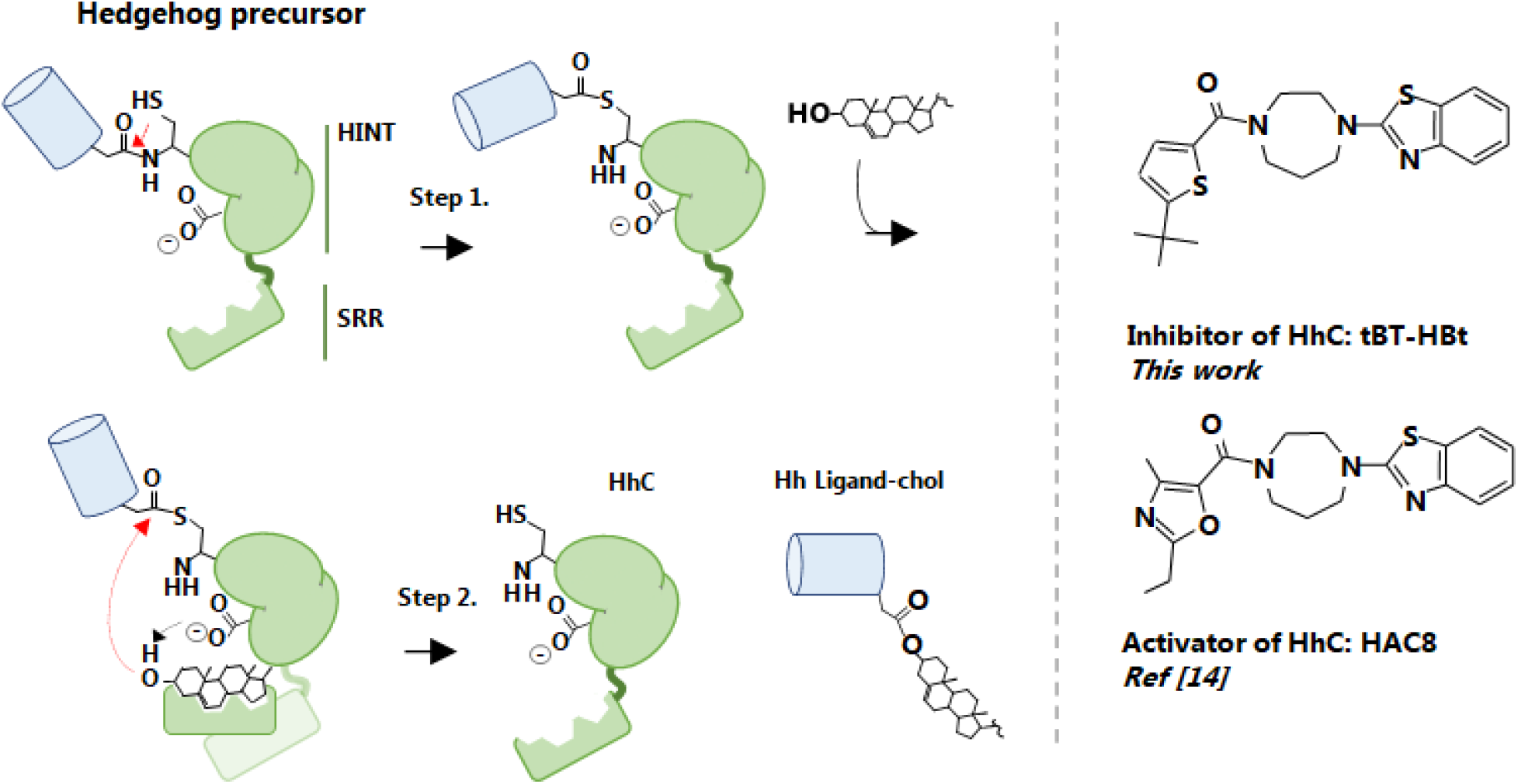
HhC functions as a dedicated self-cleaving, protein carboxy sterol ligase of the Hh ligand. The Hh precursor has two domains, an N-terminal domain (HhN, blue) and a C-terminal autoprocessing domain (HhC, green). The two subdomains of HhC are indicated, HINT (Hedgehog/Intein) and SRR (sterol recognition region). *(Left)* Hh autoprocessing has two steps. *Step*, **1** The side chain thiol of a conserved cysteine residue at position **1** of HhC attacks the amide bond of the preceding residue of HhN to form an internal thioester; *Step*, 2 Binding of cholesterol by HhC triggers transesterification to the lipid’s 3β-OH group, generating cholesterylated Hh ligand and displacing HhC. *(Right)* Representative structure of HhC inhibitor reported here and of HhC activator described previously.

We have begun pursuing selective antagonists of HhC that can phenocopy the effects of deactivating HhC mutations.^*14–16*^ Inhibitors of HhC cholesterolysis hold promise as chemical tools to understand the complex biology of Hh signaling. HhC inhibitor can also stabilize HhC’s dynamic structure ^*17*^ and enable structural and functional characterization of this unusual autoprocessing element. HhC is composed of two subdomains, the Hedgehog intein (HINT) region and the sterol recognition region (SRR). Currently only one of the two HhC subdomains (HINT) has been resolved by experimental structural methods.^*18, 19*^ Longer term, we view HhC inhibitors as novel directions for therapeutic intervention in human cancers, particularly for malignancies where existing Hh blocking drugs have lost efficacy.^*20, 21*^

Inhibiting protein autoprocessing, where the catalytic domain is tethered to its substrate, presents a special challenge. In hedgehog cholesterolysis, HhC and the substrate HhN are linked in the Hh precursor protein. There are substantial chelate-type connectivity effects between the HhC catalyst and HhN substrate that must be overcome for effective inhibition.^*22–24*^ Additional challenges arise from the single turnover nature of protein autoprocessing events like Hh cholesterolysis. Suppressing a single turnover process by an intramolecular catalyst demands inhibitors that are bound tightly. Inhibitor dissociation would permit protein autoprocessing to begin, changing the active site structure and altering inhibitor recognition. Accordingly, small molecule antagonists of Hh cholesterolysis and other forms of protein autoprocessing nearly all depend on binding irreversibly through covalent interactions.^*15, 16, 25–28*^

Here we describe an exception to the empirical “irreversibility rule” for protein autoprocessing inhibition. Screening a focused library of steroid mimetics led to the identification of noncovalent, tight-binding inhibitors of HhC cholesterolysis. We report an IC_50_ value of 300 × 10^−9^ M for the most effective inhibitor. A mixed inhibition mode is suggested by kinetic analysis and by photoaffinity labeling. Computational modeling of HhC and mutagenesis points to an allosteric binding site comprised of residues from HhC’s two subdomains. We speculate that inhibitor binding blocks conformational changes necessary for cholesterolysis. There is a striking resemblance between the inhibitors described here and a family of noncovalent, allosteric *activators* of HhC, which we described previously (**Figure 1**, *right*).^*14*^ The inhibitor/activator duality appears to be mediated by the same allosteric site on HhC, and stem from subtle differences in the structure of a heterocycle substituent on the effector.

## RESULTS AND DISCUSSION

### Library screening for HhC inhibitors

We tested 1187 steroid analogs from ChemBridge as potential inhibitors of Hh cholesterolysis using continuous light-to-dark FRET activity assay.^*14–16*^ The FRET reporter construct, C-H-Y, has HhC flanked by cyan fluorescent protein (C) and yellow fluorescent protein (Y), providing the FRET donor and acceptor, respectively. FRET is measured as the ratio of emission of 540 nm / 460 nm after excitation at 400 nm. Cholesterolysis of the engineered C-H-Y precursor protein results in the loss of FRET as the reaction products HhC-YFP (H-Y) and sterylated CFP (CFP-chol) separate (**Figure 2A**).

**Figure 2.**
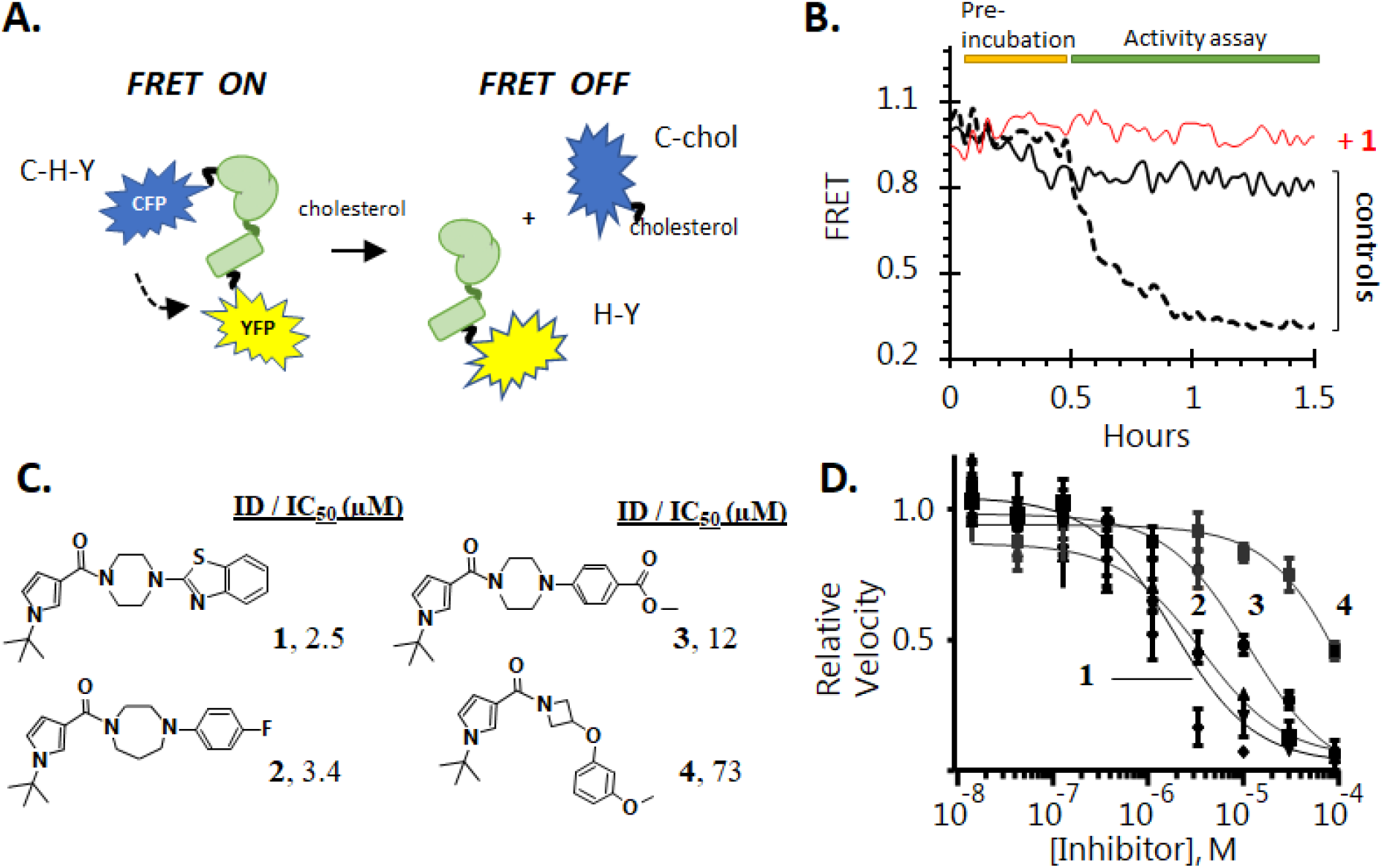
Screening assay and characterization of noncovalent HhC inhibitors. (A) Tripartite HhC construct, C−H−Y, where HhC is flanked by cyan fluorescent protein and yellow fluorescent protein is FRET active. Hh autoprocessing in the presence of cholesterol generates H-Y and C-chol, resulting in a loss of FRET signal. (B) Raw data from library screen showing inhibition by compound **1**. FRET signal from C-H-Y is plotted as a function of time, during a preincubation period (30 min) and during activity assay period (1 hr) in the presence (*black, dotted*) and absence of cholesterol (*black, solid*). Sample well with C-H-Y and 50 μM compound **1** with cholesterol (*red*) showing significant inhibition. (C) Chemical structures of screen hit compounds **1-4**, and corresponding IC_50_ values. (D) Dose Response inhibition of C-H-Y cholesterolysis by compounds **1-4** in the presence of 1.5 μM cholesterol.

The target protein in the screen was HhC from *Drosophila melanogaster*. There is 35% identity between the amino acid sequence of *Drosophila melanogaster* HhC and human HhC domains. To date, attempts to prepare recombinant human HhC protein in *E. coli* have been undermined by poor expression, insolubility, and proteolysis.

Standard reaction conditions for compound screening using C-H-Y in 96-well plates have been described.^*14*^ Briefly, wells contained Ni-NTA purified C-H-Y (100 nM) in cholesterolysis buffer (Tris, 20 mM, pH 7.1; Fos-Choline 12, 1.5 mM; EDTA, 5 mM; DTT, 1 mM) in a total volume of 100 μl. Library compounds were added from 10 mM DMSO stocks to final concentration of 50 μM. Prior to initiating the reaction with cholesterol, plates were incubated for 30 min at 30°C while FRET readings where recorded every 30 sec. Data collected during this delay were monitored to flag compounds that interfered with C-H-Y signal. Cholesterolysis was initiated by adding cholesterol from an ethanol stock (1.5 μM, final) and FRET was recoded every 30 seconds for 1.5 hours. All plates contained 8 controls wells with C-H-Y plus cholesterol, simulating no inhibition of Hh autoprocessing, and 8 wells with C-H-Y but without cholesterol, simulating 100% inhibition.

### Hit identification and validation

A total of five compounds from the library displayed inhibitory activity toward C-H-Y. In **Figure 2B** we show kinetic traces for C-H-Y in the presence of the most effective inhibitor (1) from the screen along with intraplate control samples.

Four of the five preliminary hits (1-4), available for repurchase as powders, were re-tested by dose-response assay and by end point analysis using SDS-PAGE (**Figure SI 1**). The four molecules possess similar structural components: N *tert*-butyl pyrrole, aliphatic cyclic diamine, and an aryl group (**Figure 2C**). For the dose-response assay, compounds were brought up in DMSO to 10 mM, then diluted from 100 to 0.015 μM into wells containing C-H-Y (0.1 μM), over a 9-point titration. Cholesterol was added at its KM value of 1.5 μM ^*29*^ to initiate the reaction and FRET readings were collected every 2 min. Initial velocity values were calculated as the slope of the linear phase of the FRET readings at each inhibitor concentration. Relative velocity (RV) was calculated by dividing the initial velocity with inhibitor by the initial velocity in the absence of inhibitor. Plots of the RV values as a function of log[inhibitor] (**Figure 2D**) were used to calculate IC_50_ values (Supplemental Information). IC_50_ values for **1-4** are reported as the average from ≥3 experiments resulting in the following IC_50_ values: **1**, 2 μM; **2,** 4 μM; **3** 12 μM; **4**, 180 μM. The rank order corresponds to the data obtained from the library screen.

### Kinetic evidence of mixed-type inhibition; compound **1** diminishes catalysis through effects on k_max_ and K_M_

We assayed the kinetic parameters, k_max_ and K_M_ for cholesterolysis using C-H-Y in the presence of **1** to understand the mode of inhibition. The dose response experiments above indicated an IC_50_ value of 2.0 μM. In the kinetic experiments described here, **1** was added to C-H-Y at concentrations of 0.8, 2.5, 5 and 10 μM and 25 μM, corresponding to 0.4 – 12.5 x the IC_50_ value. To those wells, we titrated substrate cholesterol at concentration range of 0.2 μM to 100 μM. In **Figure 3A**, initial cholesterolysis velocity is plotted as a function of increasing cholesterol concentration. The initial velocity data were fit initially to a standard Michaelis Menten equation (Supplemental Information). Comparison of the kinetic values +/− added **1** suggested mixed-type inhibition, where both K_M_ and k_max_ values are perturbed by an inhibitor. Calculated K_M_ values under the conditions tested were as follows: no inhibitor, 1.2 μM; with 0.8 μM **1**, 1.1 μM; with 2.5 μM **1**, 1.6 μM; and with 10 μM **1**, 2.6 μM, and with 25 μM **1**, 4.0 μM. The calculated KM values appear modestly sensitive to increasing concentrations **1**, with 2-fold increase in KM with 25-fold increase in compound **1** concentration. There was a clearer concentration dependent suppression of k_max_ with added **1**. In the absence of inhibitor, k_max_ for cholesterolysis is 1.9 × 10^−3^ sec^−1^. With 0.8 μM **1**, k_max_ is reduced to 1.3 ×10^−3^ sec^−1^ and at 2.5 μM **1**, k_max_ is reduced further to 0.9 ×10^−3^ sec^−1^. Increasing **1** to 10 and 25 μM decreases k_max_ to 0.48 x10^−3^ sec^−1^ and 0.29 ×10^−3^ sec^−1^, respectively. In summary, inhibitor **1** interferes mainly with turnover rate (6.5-fold decrease with 25-fold increase in **1**) along with minor but reproducible weakening affinity for substrate cholesterol. These results are consistent with mixed-type inhibition.

**Figure 3.**
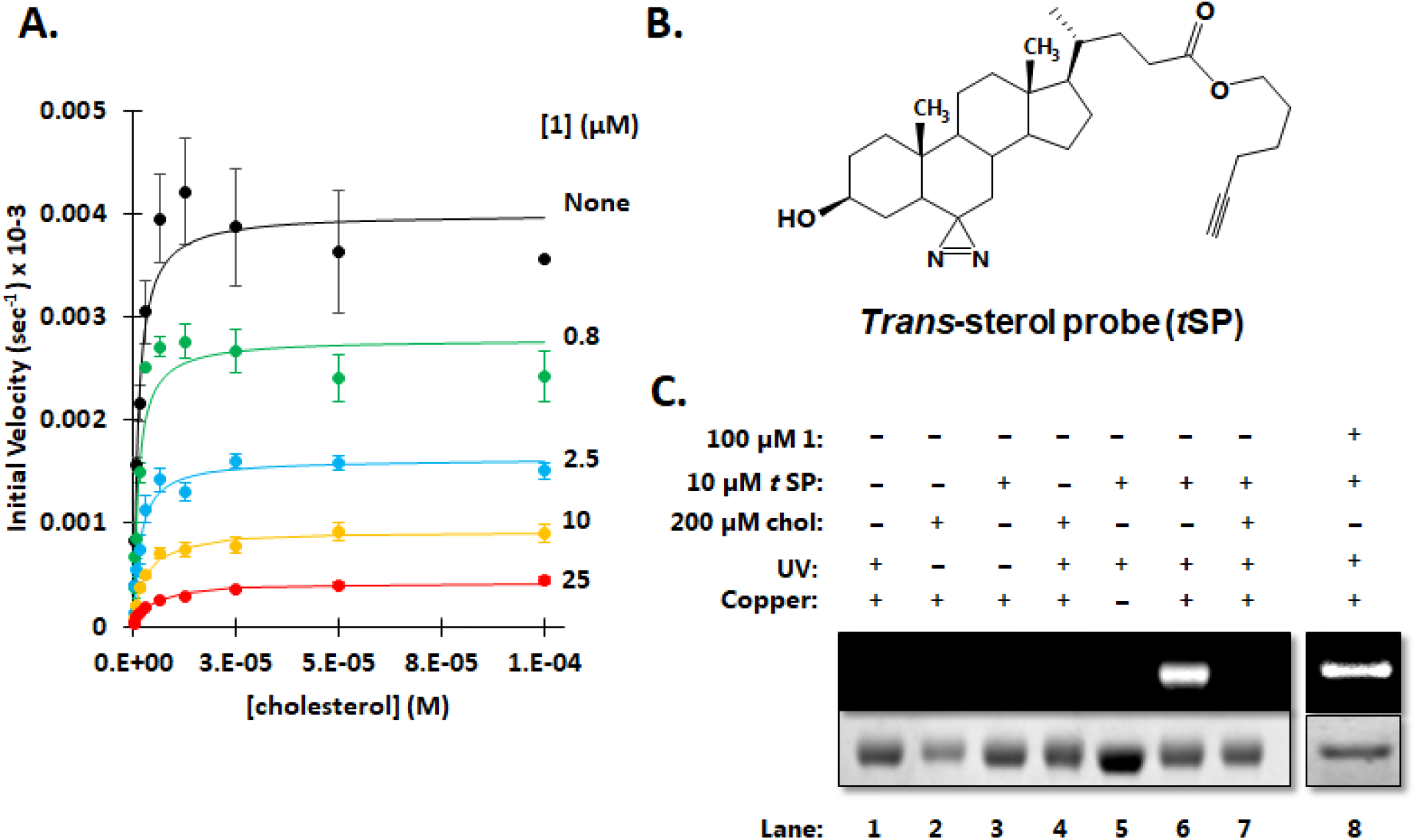
Kinetics and photolabeling suggest inhibitor **1** and substrate sterol can be bound by HhC simultaneously. (A) Mixed-type inhibition kinetics. Michaelis-Menten plot in the presence of inhibitor **1**. Initial velocity of C-H-Y cholesterolysis with indicated concentrations of compound **1**, along with no inhibitor control (*black*). (B) Photoactivatable *trans*-sterol probe (*t*SP) for HhC. (C) Photolabeling of HhC by tSP persists despite saturating concentrations of **1**. All samples contained HhC at 5 μM and 50 μM azide-linked fluorophore; other components varied as indicated. The extent of photolabeling was assessed by SDS-PAGE and fluorescence gel imaging. *(Top)* fluorescence image showing HhC-*t*SP-fluorophore adduct. *(Bottom)* Coomassie stain for total HhC protein. Controls: Lanes 1-5. Experimental: Lane 6, photolabeled HhC by *t*SP in the absence of **1**. *Lane*, 7 diminished photolabeling of HhC by *t*SP with saturating cholesterol. *Lane 8*. Persistent *t*SP photoaffinity labeling in the presence of saturating compound **1**.

### Saturating concentrations of (1) did not diminish photolabeling of HhC by diazirine-modified sterol: existence of an HhC-sterol-inhibitor complex

Mixed type inhibitors can be bound by enzymatic catalysts both in the presence and in the absence of substrate. As a complement to the above kinetic analysis, we applied photoaffinity labeling (PAL) with the *trans* sterol probe (*t*SP) to investigate the possibility of an “E-S-I” complex. *t*SP contains a photoactivatable diazirine group at position six of the sterol nucleus (**Figure 3B**).^*30*^ Alkyl diazirine groups decompose under UV light to produce N2 gas, exposing a reactive carbene or diazo intermediate.^*31*^ Covalent addition at that reactive group furnishes a chemical beacon for identifying the sterol binding sites through analytical techniques.^*17, 32, 33*^ The sterol side chain in *t*SP is also modified to include an alkyne group for click-type labeling. Despite the dual modifications, we found *t*SP was readily accepted by HhC as an alternative substrate, with K_M_ and k_max_ values of 6.8 μM and 4.2 ×10^−3^ sec^−1^ (**Figure SI 2A, B**).

Photoaffinity labeling of HhC with *t*SP was established first by click chemistry using an azide-modified fluorophore. The HhC-*t*SP-fluorophore adduct was resolved by denaturing SDS-PAGE and detected by fluorescence gel imaging. Gel mobility corresponded to the expected molecular weight (**Figure SI 2C**). To locate the *t*SP binding site in HhC, we isolated 10 μg of HhC-*t*SP adduct by reverse phase chromatography followed by MS/MS analysis. MS/MS analysis indicated *t*SP attachment to HhC peptide EQLHSSPK (HhC residues 174-181) with residues E174, P180, and K181 identified as the most probable sites of covalent photolabeling by *t*SP. These three residues map to the sterol recognition region (SRR) of HhC.

Confident in the utility of tSP as a photolabile substrate analog for HhC, we assessed the possibility of E-S-I formation by combining HHC, *t*SP and **1**. If inhibitor **1** and tSP bind at *separate* sites, photoaffinity labeling of HhC by tSP would likely be insensitive to added **1**. If inhibitor **1** and *t*SP bind at the same site, addition of **1** should block *t*SP binding by HhC and eliminate photoaffinity labeling. The results of control reactions with *t*SP and HhC are shown in lanes 1-7 of **Figure 3C**. Importantly, photolabeling of HhC by tSP could be eliminated by saturating concentrations of cholesterol (200 μM), **Figure 3C** *Lane 7*. In **Figure 3C** *Lane 8*, we show test sample containing HhC (5 μM), tSP (10 μM) and **1** (100 μM). Despite saturating amounts of **1**, the inhibitor, unlike cholesterol, did not substantially diminish photolabeling by *t*SP. The lack of, or incomplete displacement of *t*SP by saturating **1** is consistent with HhC binding of **1**at a site that is not strictly competitive with sterol. Thus, photolabeling of HhC by *t*SP in the presence of **1** is in accord with the existence of a HhC-sterol-**1** ternary complex.

### Analog testing identified nanomolar inhibitor of HhC cholesterolysis, tBT-HBt

We next prepared and tested several analogs (**Figure 4A**) as potential HhC antagonists to explore the structure-activity relationship (SAR) of inhibition. The first analog **5** incorporates the benzothiazole and N-tert butyl pyrrole groups of **1** and the homopiperazine of **2**. As an inhibitor of cholesterolysis, the molecule was improved by 2-3 fold over **1** and **2**, with an IC_50_ of 1.2 μM in the FRET assay. In analog **6** we replaced the t-butyl group of **5** by a hydrogen atom; this truncation resulted in a substantially weakened inhibitor, with an IC_50_ of 61 μM. The t-butyl group was restored in analogs **7** and **8** while the heterocycle was switched from the pyrrole to a 1,3 oxazole **7** or thiophene **8**. Compound **7** was indistinguishable from **5** as a cholesterolysis inhibitor. The thiophene **8** by contrast provided a ~ 5-fold boost in binding compared with analog **6**. Analog **8** is thereby the first nM inhibitor of HhC cholesterolysis, with an IC_50_ 300 nM. Four analogs **9-12**based on **8**were prepared next. The t-butyl group was truncated in **9**, expanded in **10** or substituted in **11-12**. Each change diminished inhibition to varying degrees, showing again the importance of the t-butyl group for inhibitor recognition by HhC. Analog **8** emerged from this panel as the optimal inhibitor, improving the IC_50_ by ~10-fold from the top hit from library screening.

**Figure 4.**
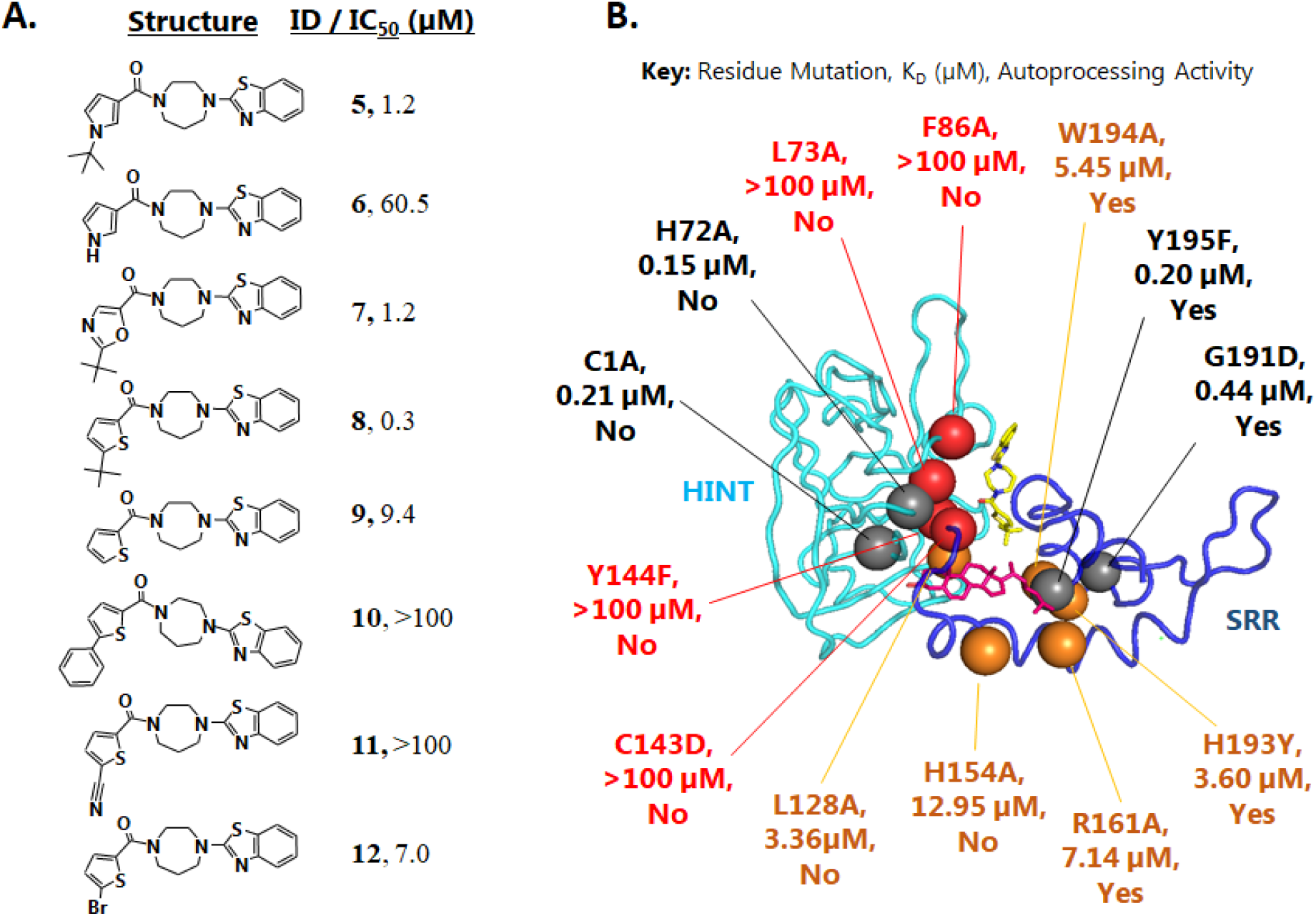
Nanomolar HhC inhibitor from analog screening and evaluation of potential inhibitor binding site by computational modeling and point mutagenesis. (A) Chemical structures of HhC inhibitor analogs, along with IC_50_ values toward cholesterolysis derived from the C-H-Y FRET assay. Compound **8** displayed sub-micromolar inhibition. (B) Proposed ternary complex model for substrate and inhibitor binding by HhC. Structure color key: inhibitor **8** or tBT-HBT, *yellow*; cholesterol, *magenta*; for HhC, the HINT subdomain, *cyan*; the SRR, *purple*. Point mutations are mapped along with the corresponding K_D_ values derived from ΔFRET titration. The mutant’s cholesterolysis activity is indicated by: yes (active) /no (not active). Mutation color key: *red*, complete loss of inhibitor binding; *orange*, 60-300 fold loss of inhibitor binding; *grey*, <10-fold loss of inhibitor binding affinity.

### Inhibitor binding by HhC is weakened by mutations in SRR and HINT subdomains of HhC

We used the nM inhibitor, **8**, hereafter called tBT-HBt, along with thirteen HhC mutants to investigate the inhibitor binding site. Our kinetic analysis above indicated mixed-type inhibition, and photolabeling experiments suggested that HhC can accommodate substrate sterol and inhibitor simultaneously. Additional support for a ternary complex, HhC-substrate-inhibitor, was obtained from a computational model of *Drosophila melanogaster* HhC, which we prepared using MODELLER ^*34*^ and the predicted human Sonic HhC protein structure ^*17*^. Virtual docking of substrate cholesterol and the tBT-HBt inhibitor into the predicted *Drosophila melanogaster* HhC structure is shown in **Figure 4A**. Single point mutations in HhC were introduced based in part on this predicted ternary structure as well as on residue conservation across HhC sequences.

The binding affinity of tBT-HBt by HhC and HhC mutants was characterized from an inhibitor dependent increase in FRET we observed with the C-H-Y reporter. By using ΔFRET to report inhibitor binding we could evaluate a greater variety of HhC mutants, including mutants that lack catalytic activity. In two previous studies we used ΔFRET from C-H-Y to monitor inhibitor binding ^*15, 16*^. In those earlier reports, we were studying covalent inhibitors and there, formation of the covalent adduct with C-H-Y caused the FRET signal to decrease. The inhibitor dependent FRET increase observed here from C-H-Y with added tBT-HBt was not apparent in C-Y, a control where HhC is replaced by a flexible peptide linker. A FRET increase was also absent with C-HINT-Y, a truncation mutant of HhC where the SRR subdomain is deleted. C-Y and C-HINT-Y were insensitive by FRET to added tBT-HBt up to 100 μM (**Figure SI 3**). The insensitivity of C-Y and C-HINT-Y to tBT-HBt suggest that this inhibitor is not bound by the two negative control constructs.

Results of our mutagenesis study are summarized in **Figure 4B**. Plots for ΔFRET as a function increasing tBT-HBt concentration for wild-type and HhC point mutants are found in **Figure SI 4**. Data were fit to a quadratic binding equation to derive an observed K_D_ value. While all point mutations we tested resulted in decreased tBT-HBt binding, we were able to group the mutational effects into three general categories: <10-fold loss of binding affinity (**Figure 4B**, grey); between 60-300 fold loss of binding affinity (**Figure 4B**, orange); abolished inhibitor binding (**Figure 4B**, red). The most severe loss in tBT-HBt binding mapped to mutations in the HINT domain. These residues (L73A, F86A, C143D, Y144F, colored red) form part of lining for the predicted tBT-HBT binding site in the computational model in **Figure 4B**. Mutations that produced a large but not total loss, 60-300 fold, of tBT-HBt binding mapped to the SRR subdomain (H154A, R161A, H193Y, W194A, colored orange). Two mutations in HINT (C1A, H72A, colored grey) and two in the SRR (G191D, Y195F, colored grey) produced small losses in tBT-HBt binding affinity. Among sites whose mutation reduced binding by more than 60 fold, only W194 is predicted to contact the inhibitor directly in the model. The compromised binding in H154A, R161A, and H193A, which are distal to the predicted binding site, lack clear explanation at this time. Collectively, the results support speculation that the inhibitor binding depends on interactions with residues in the SRR and HINT subdomains, in general accord with the modeling and docking of **Figure 4B**.

## CONCLUDING REMARKS AND INHIBITOR / ACTIVATOR DUALITY

HhC, the C-terminal half of Hedgehog precursor proteins, possesses cholesterolysis activity that is necessary to release and sterylate the adjacent Hh signaling ligand, HhN. The significance of this initiating step in the Hh signaling pathway is three-fold: the transformation appears unique to Hh family proteins; it lies upstream of all known receptor interactions; and lastly, sterylation provides a vital membrane anchor that regulates Hh ligand diffusion and physiological signaling. Small molecules that manipulate HhC autoprocessing could help unravel the complex biology of Hh signaling by providing an off switch at the origin of the Hh signaling pathway. The promise of HhC antagonists/agonists extends to therapeutics in treating sporadic tumors and congenital disorders where Hh signaling is dysregulated.

Antagonists for intramolecular catalytic systems like Hh cholesterolysis seem destined to act by covalent mechanisms. Published examples of autoprocessing inhibitors are indeed dominated by irreversible protein-inhibitor interactions, and the few examples of noncovalent autoprocessing inhibitors show conspicuously modest binding affinity.

The findings presented here represent an important exception to the “irreversibility rule” for protein autoprocessing inhibition. We speculate that the tightest binding inhibitor, tBT-HBT (**8**), traps the SRR and HINT subdomains of HhC in a conformation that is capable of binding cholesterol but incapable of catalyzing cholesterolysis. All data collected to date suggest an allosteric binding site in HhC comprised of HINT and SRR residues. To our knowledge, tBT-HBT is the tightest binding noncovalent inhibitor of HhC cholesterolysis, and it appears to be unrivaled among reported noncovalent inhibitors of protein autoprocessing in general.

In an earlier report we described reversibly bound small molecules targeting HhC that *activate* cholesterol-*independent* autoprocessing, a nonnative paracatalytic reaction. The structures of the best HhC activator, HAC8, and most potent inhibitor described here, tBT-HBT, are strikingly similar (**Figure 1**). The compounds emerged from the same screen and, based on competitive-type kinetics between HAC8 and tBT-HBT (**Figure SI 5**), the activators and inhibitors seem to share a binding site on HhC. Thus, depending on the structure of the bound effector, HhC can accelerate cholesterol-independent autoproteolysis, as with HAC8, or suppress the rate of cholesterolysis, as with tBT-HBT. A single allosteric site that can toggle between activation and inhibition depending on the effector is intriguing and finds precedent in selected metabolic and signaling enzymes.^*35, 36*^ Despite descriptions of protein autoprocessing domains as primitive or proto-zymes, left over from an earlier age, we continue to find close parallels between their mechanisms and extant enzymes. Moreover, the notion that protein autoprocessing domains like HhC and inteins represent intractable targets for noncovalent inhibition should be discarded.

## ACKNOWLEDGEMENTS

We acknowledge generous support from the National Cancer Institute (CA206592) and the National Heart, Lung, and Blood Institute (HL067773). We thank Callahan lab members Xiaoyu Zhang and Dan Ciulla for help and encouragement and Dr. Juergen Schulte for expert advice with NMR.

