## Supplemental Information for "Nanomolar, noncovalent antagonism of hedgehog cholesterolysis: exception to the “irreversibility rule” for protein autoprocessing inhibition"

Table of Contents S1

| | | |
|---|---|---|
| Supporting Figure 1 | Inhibition of Cholesterol processing by SDS-PAGE | S3 |
| Supporting Figure 2 | Analysis of *t*SP as a substrate and probe for HhC | S4 |
| Supporting Figure 3 | Compound 1 ΔFRET controls | S5 |
| Supporting Figure 4 | ΔFRET Point Mutant Assay | S6 |
| Supporting Figure 5 | HAC8 and Compound 1 competition assay | S7 |
| Supporting Figure 6 | Compound 1 Lineweaver burke plot | S8 |
| Supporting Figure 7 | IC50 Plots of all synthesized compounds | S9 |

| | | |
|---|---|---|
| Chemicals, Solvents, Purification Buffers | | S10-S11 |
| Chemical Library | | S11 |
| Generation of Plasmid Constructs | | S11 |
| Expression and Purification of Protein | | S11 |
| HhC Inhibitor screening | | S11 |
| Analysis of Cholesterolysis Kinetics | | S12 |
| Photoaffinity Labeling | | S13 |
| Synthesis of Inhibitor Analogs | | S14 |
| HhC Computational Modelling | | S16 |
| NMR Compounds **5-12** | 1H, 13C spectra | S18-33 |
| LCMS Compounds **5-12** | LC Data with Peak MS | S34 |
| References | | S35 |

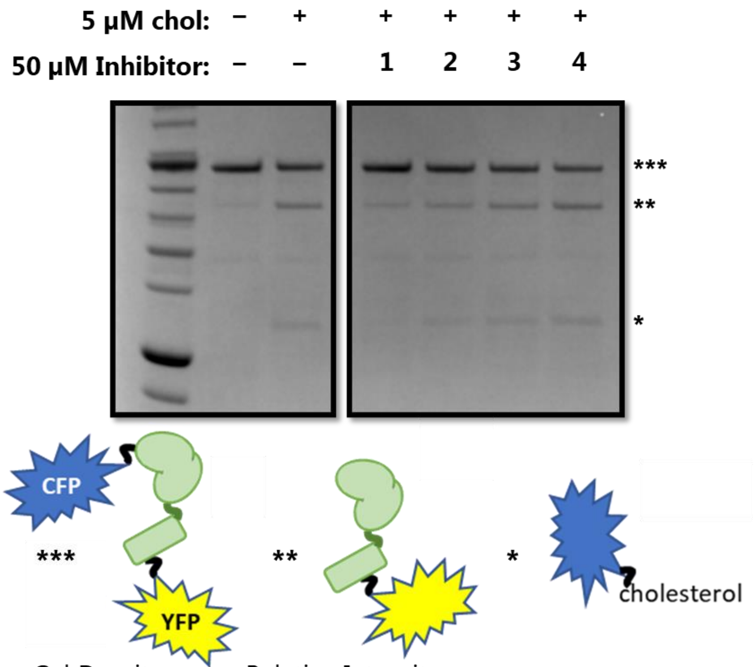


Gel Densitometry, Relative Intensity

| Band identity | No Sterol | + Chol | Inhib 1 | Inhib 2 | Inhib 3 | Inhib 4 |
|---|---|---|---|---|---|---|
| Precursor | **1.0** | 0.6 | 0.9 | 0.7 | 0.6 | 0.5 |
| HhC-YFP | 0.3 | **1.0** | 0.4 | 0.6 | 0.9 | 1.2 |
| CFP-Chol | 0.1 | **1.0** | 0.3 | 0.5 | 0.8 | 1.1 |

Figure SI 1: C-H-Y cholesterolysis activity by SDS-PAGE in the presence of screen hits **1-4** confirm compound **1** is the most effective hit. *(Top, Left panel)* Control reactions of 0.3 µM C-H-Y in presence or absence of 5 µM cholesterol. *(Top, Right panel)* Reactions with 5 µM cholesterol after 30-minute incubation with 50 µM of **1-4**. *(Bottom)* Densitometry of precursor and product bands.

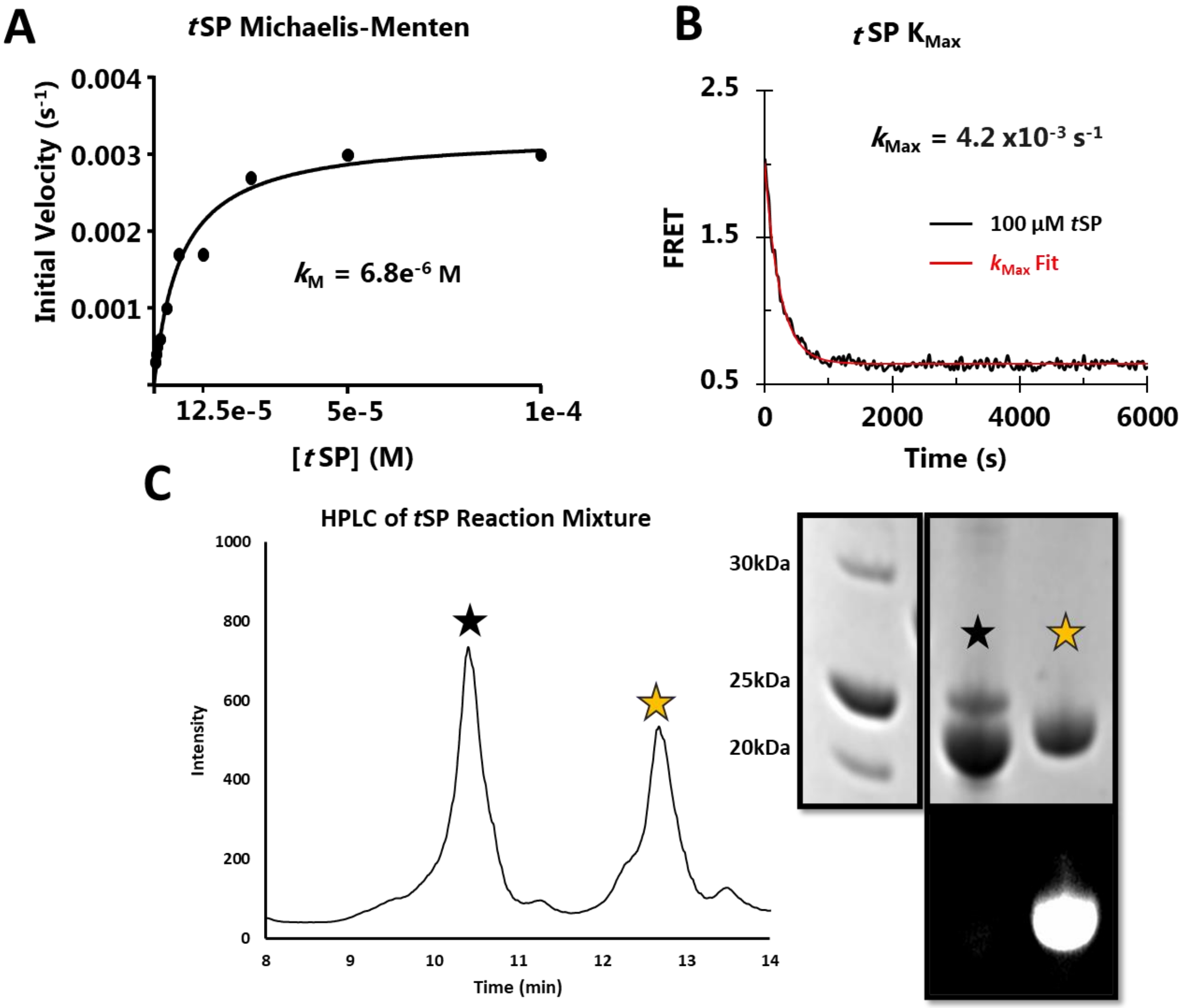


Figure SI 2: *t*SP is a substrate analog and photoaffinity label for HhC. (A) Michaelis-Menten plot of initial reaction velocity as a function of *t*SP concentration. Data were fit to a hyperbolic equation with the indicated $k_{MAX}$ and $K_M$ values. (B) Kinetic trace for C-H-Y reaction with 100 µM *t*SP reaction. Data trace was fit to a first order exponential to determine $k_{Max}$. (C) *(Left)* Reverse phase HPLC separation *t*SP-photolabeled HhC (gold star). *(Right)* SDS-PAGE confirmation of photolabeled HhC. HPLC fractions containing unlabeled (black star) and photolabeled HhC (gold star) were reacted with azide-modified fluorophore, separated by SDS-PAGE and visualized with Coomassie staining (top), and by UV imaging (bottom).

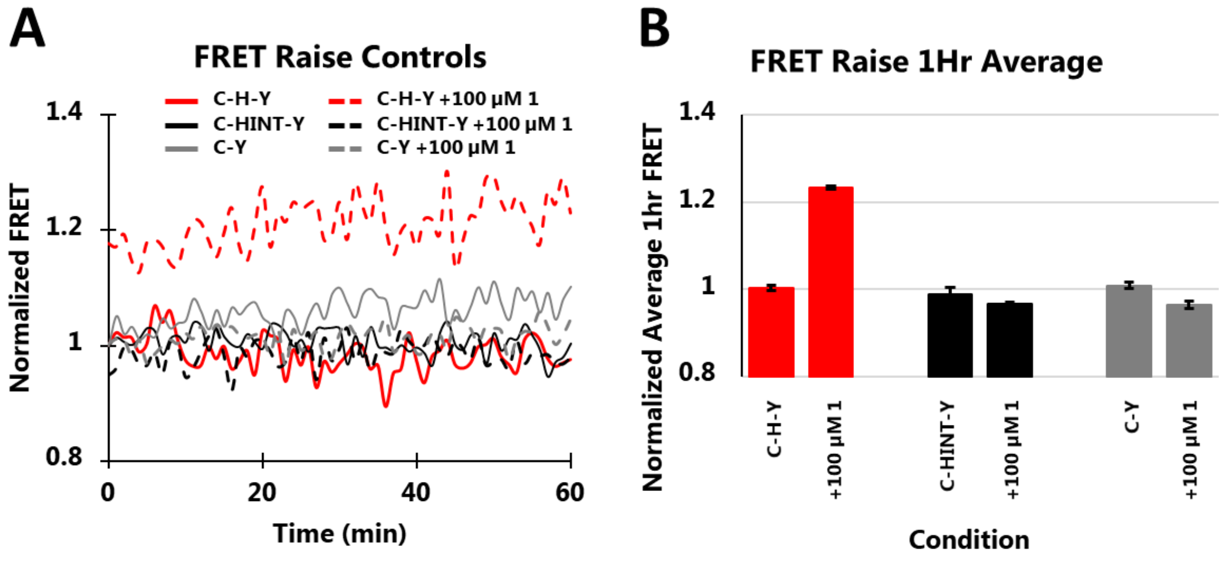


Figure SI 3: Compound **1** increase the FRET of C-H-Y not negative control constructs, C-Y and C-HINT-Y. Average FRET change after 1 hour incubation in the absence and presence of **1** (100 µM) for constructs, C-H-Y, C-HINT-Y and C-Y. Samples lacking **1** were normalized to FRET signal of unity (n=6 for each construct). (A) Full time course. (B) Data at assay end point.

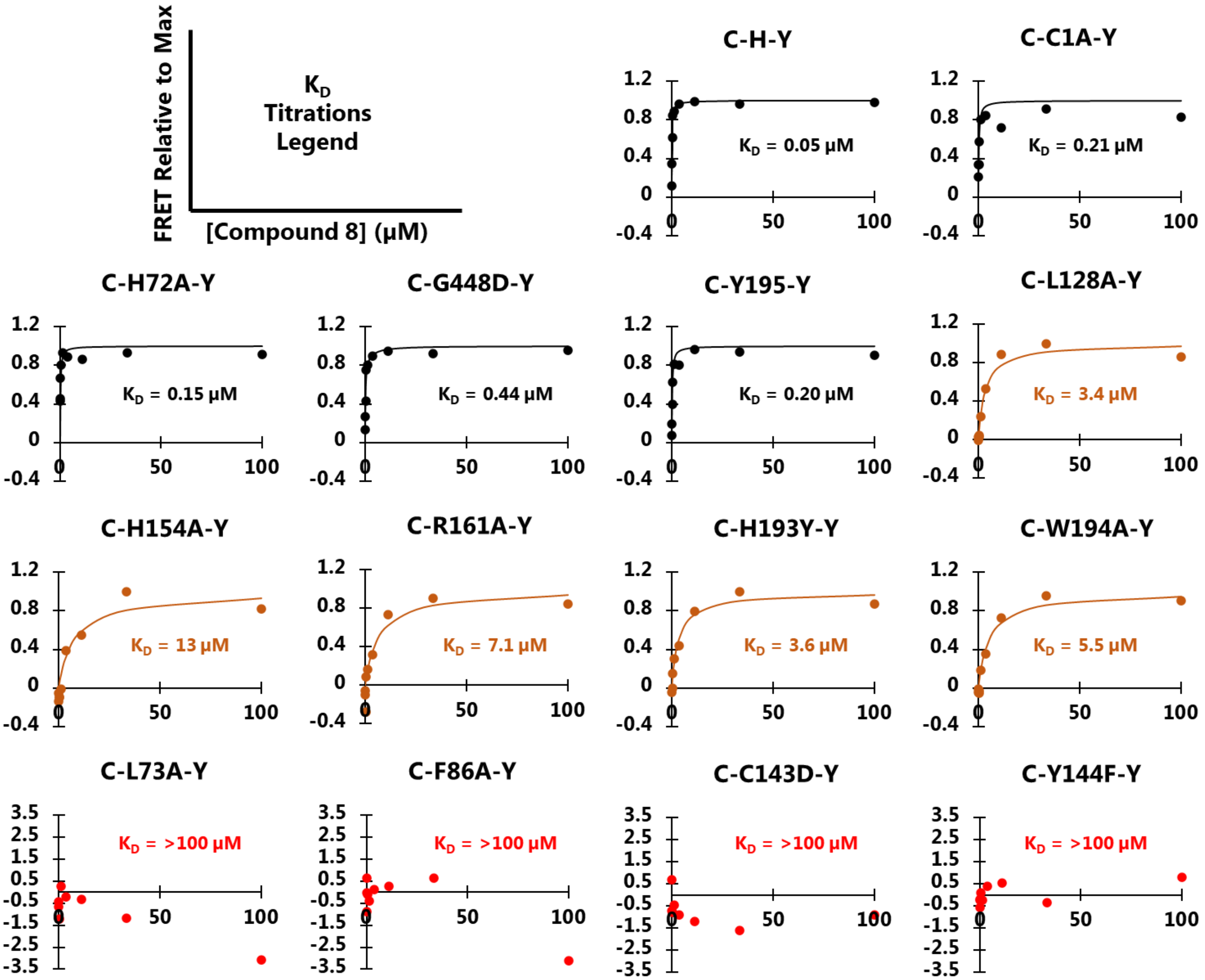


Figure SI 4: ΔFRET Inhibitor binding assay shows wide range of affinity with HhC point mutants. ΔFRET is plotted as a function of increasing compound **8** concentration. Curves represent binding isotherms for the indicated $K_D$ values except for mutants in last row (*red*) which did not appear to bind **8**.

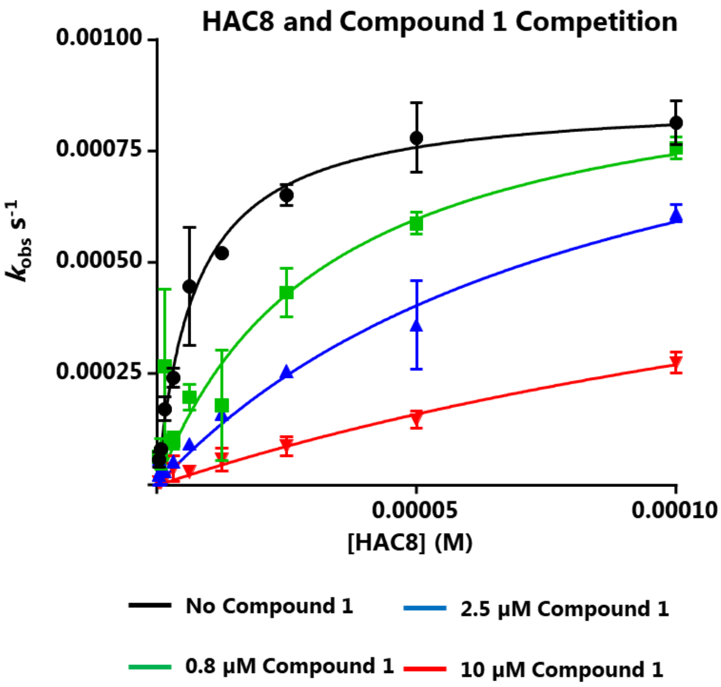


| [Compound 1] (µM) | HAC8 $AC_{50}$ (µM) | Fold Diff | HAC8 $k_{Max}$ ($s^{-1}$ x$10^{-4}$) | Fold Diff |
|---|---|---|---|---|
| 0 | 7.2 | 1.0 | 9.5 | 1.0 |
| 0.8 | 32.3 | 4.5 | 8.6 | 1.1 |
| 2.5 | 87 | 12 | 6.1 | 1.6 |
| 10 | 230 | 32 | 2.8 | 3.5 |

Figure SI 5: HAC8 and compound 1 show competitive binding kinetics. *(Left)* Michaelis-Menten plot of HAC8 titration in presence of 0.8 µM, 2.5 µM or 10 µM Compound 1. *(Right)* A table of $AC_{50}$ and $k_{Max}$ values calculated from the Michaelis Menten plot.

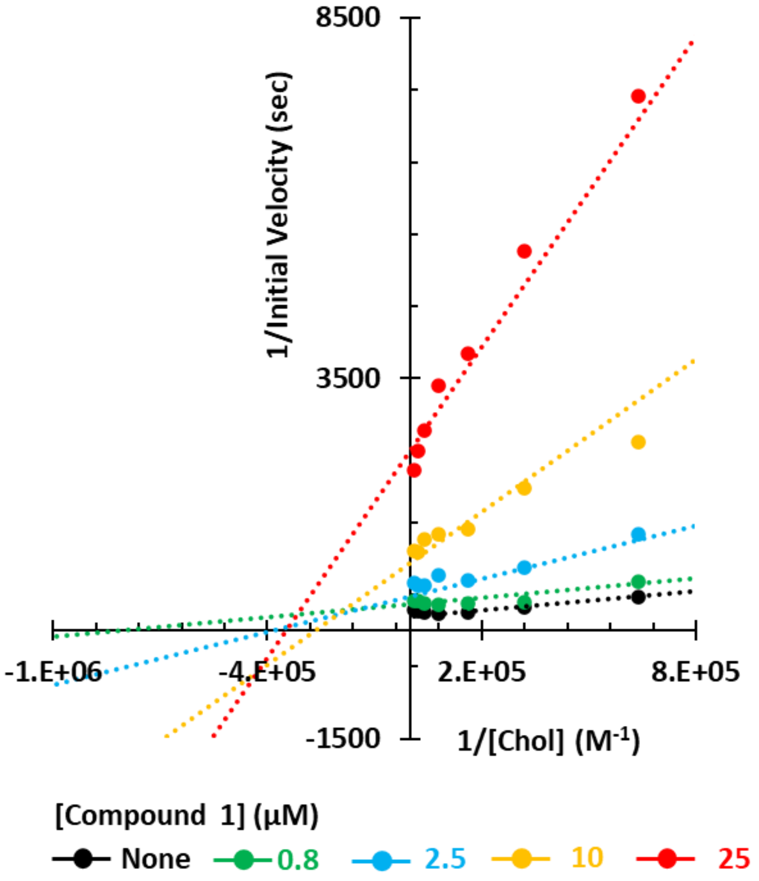


Figure SI 6: Compound **1** Lineweaver burke plot. Double reciprocal plot of titrated cholesterol with compound **1** at four fixed concentrations, against uninhibited control.

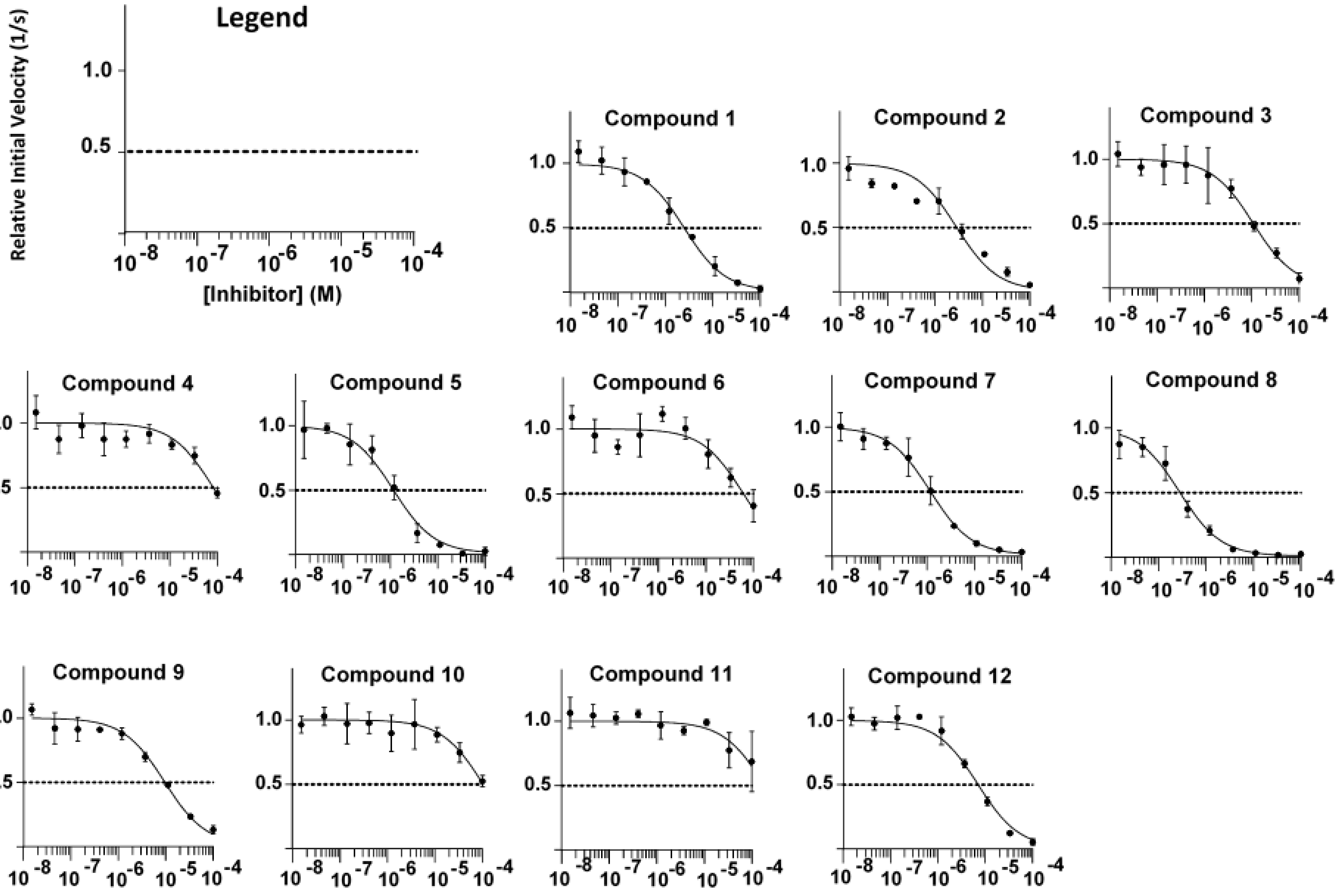


**Figure SI 7:** $IC_{50}$ Plots of all compounds based on the C-H-Y activity assay.

### Materials and Methods

**Chemicals and Solvents**: The following reagents and solvents that were used for synthesis of compounds and subsequent reactions in this manuscript were purchased from commercial vendors:

Acros Organics: silica gel, Imidazole

Alfa Aesar: Homopiperazine, 2-chlorobenzothiazole, 4-(Dimethylamino)pyridine (DMAP), L-Ascorbic Acid, 5-Bromo-2-thiophenecarboxylic acid

Anatrace: n-Dodecylphosphocholine (Fos-12)

BLD Pharm: sodium thiophene-2-carboxylate

Cambridge Isotopes: d-chloroform, d-methanol

ChemBridge, Hit2Lead: Inhibitor Compounds 1-4, 1-Tert-butyl-1H-pyrrole-3-carboxylic acid,

Click Chemistry tools: OG 488 Azide fluorophore, Tris(benzyltriazolylmethyl)amine (THPTA)

Corning: 96 Well Black Polystyrene Microplate （#3650）

Cytiva: HisSpinTrap Ni Sepharose High Performance spin columns

Enamine: 2-tert-butyl-1,3-oxazole-5-carboxylic acid, 5-cyanothiophene-2-carboxylic acid, 5-phenylthiophene-2-carboxylic acid

Fischer Scientific: dichloromethane, ethyl acetate, glycerol, tris(2-carboxyethyl)phosphine (TCEP), Tris Base

G Bioscience: dimethylformamide (DMF)

GE Healthcare: Tetramethylethylenediamine (TEMED)

InVitria: Lysozyme

Invitrogen – 40% Acrylamide (29:1 mono:bis)

JT Baker: methanol, sodium sulfate, copper sulfate

MP Biomedicals: Iodoacetamide

Millipore Sigma: Triton X-100

New England Biolabs: All restriction enzymes and buffers, DNA ligase and buffers, and all competent cells, plasmid miniprep kits

Oakwood Chemical: 5-Tert-butylthiophene-2-carboxylic acid

Qiagen: Filtered Ni-NTA spin columns

Sigma-Aldrich: 1-ethyl-3-(3-dimethylaminopropyl) carbodiimide hydrochloride (EDC), 1,4-Dithiothreitol (DTT), β-Mercaptoethanol (BME), L-Arabinose, 1H-pyrrole-3-carboxylic acid,

Thermo Fisher Scientific: All antibiotics, Isopropyl β-D-1- thiogalactopyranoside (IPTG), LB Broth Miller (Granulated), Dimethyl sulfoxide (DMSO), LB agar (miller), Coomassie Brilliant Blue G-250, Ammonium Persulfate,

VWR: Acetic Acid Glacial

**Purification Buffers:**

- Bacterial Cell Lysis buffer: 0.5 % Triton X-100, 0.05 M K2HPO4, 0.4 M NaCl, 0.1 M KCl, 10 % glycerol, 0.01 M imidazole, pH=7.3.
- Ni-NTA Bind buffer: 1 M NaCl, 0.04 M Na2HPO4, 0.01 M imidazole, 20 % glycerol, pH=7.5.
- Ni-NTA Wash buffer: 1 M NaCl, 0.04 M Na2HPO4, 0.075 M imidazole, 20 % glycerol, pH=7.5.
- Ni-NTA Elution buffer: 0.02 M Na2HPO4, 0.5 M NaCl, 0.5 M imidazole, 10% glycerol, pH=7.3.

**Chemical Library:** Compounds used in the FRET chemical screen were purchased from ChemBridge and stored at -80 degrees Celsius as 10 mM stocks dissolved in DMSO.

**Generation of Plasmid Constructs:** C-H-Y, C-Y, C-C1A-Y, and C-C143D-Y FRET constructs were cloned, characterized, and expressed as described in previous work[1-3]. Synthetic genes for HhC point mutant FRET constructs C-HINT-Y, C-H72A-Y, C-L73A-Y, C-F86A-Y, C-L128A-Y, C-Y144F-Y, C-H154A-Y, C-R161A-Y, C-G191D-Y, C-H193Y-Y, C-W194A-Y, and C-Y195F-Y were cloned using enzymes XhoI and PstI into bacterial expression plasmid pBAD33 that previously contains a C-I-Y construct gene, where "I" is a 400bp placeholder gene corresponding to a DNA E intein [4]. Following ligation with T4 polymerase, plasmids were transformed into NEB5α high efficiency competent *E. coli* cells and selected for by plating cells onto agar containing 50 µg/ml of chloramphenicol. Colonies were grown and miniprepped, and the plasmid identity was confirmed by cutting out the inserted gene with restriction enzymes XhoI and PstI. Miniprepped plasmid was then used to transform competent LMG-194 *E. coli* cells for protein expression.

**Expression and Purification of Protein:** LMG-194 *E. coli* cells containing plasmid of interest were grown from single colony in 50ml of LB media at 37 degrees Celsius and shaking 230rpm, until reaching an optical density at 600nm (OD600) of 0.5. Cultures were then induced with 100mg L-Arabinose per 50ml culture and set at 16 degrees Celsius shaking at 190 rpm for 18-24 hours. Cells were then pelleted by spinning the culture in a centrifuge at 10,000xg for 5 mins, and decanting supernatant media. Cell pellets were resuspended in 3ml of ice-cold lysis buffer with 1mg/ml lysozyme, vortexed, and sonicated. Soluble protein containing the construct of interest was separated from insoluble protein in a centrifuge by spinning at 12,000xg for 30 minutes. The construct was purified from the soluble fraction by use of Cytiva NiNTA spin columns, with the above listed bind, wash, and elution buffers. Eluted protein is analyzed for purity by 12% acrylamide SDS PAGE, and concentration of the elution is quantified by gel densitometry using ImageJ software.

**HhC activity screening:** Protein FRET Assays were monitored in 96-well plates using a BioTek Synergy H1 plate reader. Reaction conditions include FRET protein of interest (1x10-7 M) in Bis-Tris buffer (0.02 M, pH 7.1) with ethylenediaminetetraacetic acid (EDTA, 0.005 M), Sodium Chloride (NaCl, 0.1 M), n-Dodecylphosphocholine (Fos-12, 0.0015 M) and Tris(2-

carboxyethyl)phosphine (TCEP, 0.005 M). Protein and potential inhibitors were incubated for 30 minutes at 30°C at a volume of 100 µl. HhC autoprocessing was then induced by addition of the cholesterol dissolved in ethanol to a final concentration of 1.5 µM (4% ethanol v/v). FRET activity was monitored at 90 second intervals by exciting the assay at 400nm (CFP excitation wavelength) and calculating a ratio of 540nm (YFP emission from intact precursor) over 460nm (CFP emission from processed precursor), resulting in loss of FRET with successful cholesterol reaction. Cholesterol processing activity was also displayed by SDS-PAGE, evident by reduction of the ~80kDA precursor band, and appearance of two novel bands, a cholesterylated CFP (23kDa) and HhC-YFP (56 kDa) bands. A Bio-Rad Gel Doc EZ system was used to record gel images for both UV imaging and coomassie staining, and the images were subsequently quantified by ImageJ.

**Analysis of FRET Kinetics**:

$IC_{50}$ – Inhibitor was titrated into 0.1 µM protein using 1 to 3 dilutions from a 100 µM starting concentration into activity screening wells containing a set concentration of cholesterol at its previously determined $K_M$ value [5] of 1.5 µM. Relative Initial velocity was determined by calculating the initial rate of FRET loss upon addition of cholesterol of an inhibited condition and dividing it by the initial rate of FRET loss of an uninhibited condition. Relative initial velocity (RV) was plotted as a function of the log of inhibitor concentration (X) in triplicate using GraphPad which adapts the following equation:

$$\mathbf{RV = Bottom + (Top\text{-}Bottom)/(1+(X/IC_{50}))}$$

Top and bottom values were constrained to 1 and 0.01 respectively, and graphs were plotted with a Log scale on the X-axis.

$K_M$ – Cholesterol was titrated into activity screening wells containing a set concentration of inhibitor. Values for cholesterol binding affinity were derived using a Michaelis Menten graph of initial rate of FRET loss upon addition of cholesterol ($V_O$) plotted as a function of cholesterol concentration, resulting in the equation:

$$\mathbf{V_O/V_{max}=[cholesterol]/( [cholesterol]+ K_M ).}$$

Analysis of triplicate experiments and $k_M$ value was done using GraphPad software.

$k_{Max}$ – Values for first order rate of cholesterol processing were calculated by fitting the entire FRET kinetic trace of the cholesterol reaction to the following equation:

$$\mathbf{FRET=A*e^{(-kmax)*t}+C}$$

Error in the first order curve fit was reduced by calculation of square error between the FRET trace and calculated decay, and then error was reduced by use of the Solver function in Excel to refine the fit.

Mode of Inhibition Assays – Substrate (cholesterol or HAC8) was titrated into 0.1 µM protein using 1 to 2 dilutions from a 100 µM starting concentration into activity screening wells containing a set concentration of Inhibitor compound 1, at either 0.8 µM, 2.5 µM, 10 µM, or 25 µM. For assays using cholesterol, binding affinity were derived using a Michaelis Menten graph of initial rate of FRET loss upon addition of cholesterol ($V_O$) plotted as a function of cholesterol concentration, using the equation:

$$V_O/V_{max}=[cholesterol]/( [cholesterol]+ K_M).$$

For assays using HAC8 which can turn over multiple C-H-Y molecules, $k_{obs}$ values were calculated by fitting the entire trace to a first order decay curve and were plotted as a function of HAC concentration. The resulting data were fit to a modified Michaelis Menten equation below. Values for $k_{Max}$ ($sec^{-1}$) and the $AC_{50}$ (µM) were determined by nonlinear regression using Excel Solver.

$$k_{obs} = (k_{Max} *[HAC8])/(AC50 + [HAC8])$$

ΔFRET Assay – Inhibitor compound 8 was titrated into 0.1 µM C-H-Y protein. Average FRET raise over 1 hour was collected in triplicate for each condition and plotted versus inhibitor concentration using a modified quadratic equation for tight inhibitor binding, where [E] and $K_D$ represent the precursor concentration and apparent inhibition constant respectively, and $\Delta FRET_{Max}$ represents the maximum change in FRET from addition of inhibitor. Data was fit to the below equation and error was reduced by use of the Solver function in Excel to refine the fit.

$$\Delta FRET_{obs} = \Delta FRET_{Max} * \frac{(([E]+[I]+[K_D])-\sqrt{([E]+[I]+[K_D])-4*[E][I]}}{2[E]}$$

**Photoaffinity Labeling**

Preparation of HhC for Labeling Experiments – A BL21DE3 cell line transformed with ampicillin resistant pet45b plasmid containing synthetic construct 6xHIS-HhC is grown in 40 milliliters of LB Broth containing ampicillin at 37 degrees Celsius overnight. The 40ml cultures are diluted into 2 Liters of LB, grown to an optical density at 600nm (OD600) of 0.5, and then expressed with 0.5 mM IPTG for 20 hours at 16 degrees Celsius. Cell pellets were resuspended in 40ml total of ice-cold lysis buffer with 1 mg/ml lysozyme, vortexed, and sonicated. Soluble protein containing the construct of interest was separated from insoluble protein in a centrifuge by spinning at 12,000xg for 30 minutes. The 6xHIS-HhC construct is then purified from the soluble fraction by nickel affinity chromatography and cleaved of the N-terminal domain with 200 mM beta-mercaptoethanol at 16 degrees for 24 hours. 10 ml aliquots of the reaction mixture are added to dialysis tubing and dialyzed into 500ml of 1x PBS for 12 hours, twice. The dialyzed reaction mixture was then added to a Qiagen filtered nickel column that has been equilibrated with 10mM Imidazole. Cleaved and concentrated HhC containing no 6xHIS tag is collected in a 75mM Imidazole wash from the column and used for subsequent labeling experiments.

Sterol *t*SP Labeling – Reaction mixture was composed of 49 microliter aliquots of HhC (5x10-6 M) with no N-terminal domain in Bis-Tris buffer (0.02 M, pH 7.1) with sodium chloride (NaCl, 0.1 M), n-Dodecylphosphocholine (Fos-12, 0.0015 M) and L-Ascorbic acid (0.005 M). Inhibitor compound 1 was added in DMSO to the reaction mixtures to achieve a final concentration of 100µM inhibitor, resulting in 2% v/v DMSO. Cholesterol was added in ethanol to the reaction mixtures to reach a final concentration of 200 µM, resulting in 2% v/v ethanol. After these additions, all conditions were incubated for 30 minutes at room temperature, followed by addition of the *trans*-sterol probe (*t*SP) in ethanol to a final concentration of 10 µM, reaching a final ethanol concentration of 4% (v/v). The reaction was incubated at room temperature for an additional 30 minutes, and then irradiated with UV light for 5 minutes using a mercury UV lamp, with the sample in a quartz cuvette at a distance of 8 cm from the source. Samples were then denatured by addition of 18 mg (6 M) urea. A premixed copper sulfate and THPTA solution at a molar ratio of 1:5 was added to reaction mixtures to final concentrations of 1 mM Copper sulfate and 5 mM THPTA, followed by addition of OG 488 Azide fluorophore, Aminoguanidine HCL, and L-Ascorbic Acid to final concentrations of 50 µM, 5 mM and 5 mM respectively. The reaction mixture incubated at 30 degrees for 12 hours. 20 µl samples were reduced with load dye containing DTT, boiled, and loaded onto a 12% acrylamide SDS PAGE gel. A Bio-Rad Gel Doc EZ system was used to record gel images for both UV imaging and coomassie staining.

Identification of Modified Residues – Cleaved HhC protein labeled with *t*SP was injected over HPLC using a reverse-phase C4 Jupiter column, with the following buffer gradient. Line A – Water+0.1% TFA, Line B – Acetonitrile+0.1%TFA. 25%B increasing to 55% B over 20 minutes, with a flow rate of 0.8ml/min. Collected peaks were combined, evaporated to dryness, and resuspended in 20 µl of deionized water. All resuspended collections were subject to a premixed copper sulfate and THPTA solution at a molar ratio of 1:5 was added to reaction mixtures to final concentrations of 1 mM Copper sulfate and 5 mM THPTA, followed by addition of OG 488 Azide fluorophore, Aminoguanidine HCL, and L-Ascorbic Acid to final concentrations of 50 µM, 5 mM and 5 mM respectively. The reaction mixture incubated at 30 degrees for 12 hours, and then boiled with DTT load dye before analyzing by a 12% SDS PAGE gel. A Bio-Rad Gel Doc EZ system was used to record gel images for both UV imaging and coomassie staining, to confirm labeled collected peaks. The labeled peaks were then collected in subsequent injections, combined until a mass of 10 µg total was collected and confirmed by densitometry from SDS-PAGE. The 10 µg was subject to proteolytic digest and analyzed by MS/MS.

**Synthesis of Novel Compounds**:

The Homopiperazine-Benzothiazole amine fragment was synthesized from Homopiperazine and 2-chlorobenzothiazole as previously described [6].

(a) **Compound 5**: 25.1 mg (.15 mMol, 1.5 equivalents) of 1-Tert-butyl-1H-pyrrole-3-carboxylic acid, 2.44 mg (0.02 mMol, 0.2 equivalents) of 4-(Dimethylamino)pyridine (DMAP), and 38.34 mg (0.2 mMol, 2 equivalents) of N-(3-Dimethylaminopropyl)-N'-ethylcarbodiimide hydrochloride (EDC) were mixed in 4 ml of dichloromethane, and incubated for 30 minutes at room temperature stirring. 23.3 mg (0.1 mMol, 1 equivalent) of Homopiperazine-benzothiazole amine was dissolved in dichloromethane and added to the aforementioned mixture and allowed to react for 4 hours. The solution was aqueous extracted, dried with anhydrous sodium sulfate, and the product was separated via silica gel flash chromatography with isocratic 1:1 dichloromethane:Ethyl acetate as the solvent. Product was isolated in 90% yield.

(b) **<u>Compound 6</u>**: 11.1mg (0.1 mMol, 1 equivalent) of 1H-pyrrole-3-carboxylic acid, 2.44 mg (0.02 mMol, 0.2 equivalents) of 4-(Dimethylamino)pyridine, and 38.34 mg (0.2 mMol, 2 equivalents) of N-(3-Dimethylaminopropyl)-N′-ethylcarbodiimide hydrochloride were mixed in 4 ml of 50% Acetone/50% dichloromethane (v/v), and incubated for 30 minutes at room temperature stirring. 23.3 mg (0.1 mMol, 1 equivalent) of Homopiperazine-benzothiazole amine was dissolved in dichloromethane and added to the aforementioned mixture and allowed to react for 4 hours. The solution was aqueous extracted, dried with anhydrous sodium sulfate, and the product was separated via silica gel flash chromatography with isocratic 9:1 Ethylacetate:ethanol. Product was isolated in 66% yield.

(c) **<u>Compound 7</u>**: 16.9 mg (0.1 mMol, 1 equivalent) of 2-tert-butyl-1,3-oxazole-5-carboxylic acid, 2.44 mg (0.02 mMol, 0.2 equivalents) of 4-(Dimethylamino)pyridine, and 38.34 mg (0.2 mMol, 2 equivalents) of N-(3-Dimethylaminopropyl)-N′-ethylcarbodiimide hydrochloride were mixed in 4 ml of dichloromethane, and incubated for 30 minutes at room temperature stirring. 23.3 mg (0.1 mMol, 1 equivalent) of Homopiperazine-benzothiazole amine was dissolved in dichloromethane and added to the aforementioned mixture and allowed to react for 4 hours. The solution was aqueous extracted, dried with anhydrous sodium sulfate, and the product was separated via silica gel flash chromatography with isocratic 9:1 Ethylacetate:ethanol. Product was isolated in 68% yield.

(d) **<u>Compound 8</u>**: 18.4 mg (0.1 mMol, 1 equivalent) of 5-Tert-butylthiophene-2-carboxylic acid, 2.44 mg (0.02 mMol, 0.2 equivalents) of 4-(Dimethylamino)pyridine, and 38.34mg (0.2 mMol, 2 equivalents) of N-(3-Dimethylaminopropyl)-N′-ethylcarbodiimide hydrochloride were mixed in dichloromethane, and incubated for 30 minutes at room temperature stirring. 23.3 mg (0.1 mMol, 1 equivalent) of Homopiperazine-benzothiazole amine was dissolved in dichloromethane and added to the aforementioned mixture and allowed to react for 1-4 hours. The solution was aqueous extracted, dried with sodium sulfate, and the product was separated via silica gel flash chromatography with isocratic 9:1 Ethylacetate:Ethanol as the solvent. Product was isolated in 44% yield.

(e) **<u>Compound 9</u>**: 15mg (0.1 mMol, 1 equivalent) of sodium thiophene-2-carboxylate, 2.44 mg (0.02 mMol, 0.2 equivalents) of 4-(Dimethylamino)pyridine, and 38.34 mg (0.2 mMol, 2 equivalents) of N-(3-Dimethylaminopropyl)-N′-ethylcarbodiimide hydrochloride were mixed in 4ml of dichloromethane, and incubated for 30 minutes at room temperature stirring. 23.3 mg (0.1 mMol, 1 equivalent) of Homopiperazine-benzothiazole amine was dissolved in dichloromethane and added to the aforementioned mixture and allowed to react for 4 hours. The solution was aqueous extracted, dried with anhydrous sodium sulfate, and the product was separated via silica gel flash chromatography with isocratic 9:1 dichloromethane:Methanol. Product was isolated in 80% yield.

(f) **Compound 10**: 20.4mg (0.1 mMol, 1 equivalent) of 5-phenylthiophene-2-carboxylic acid, 2.44 mg (0.02 mMol, 0.2 equivalents) of 4-(Dimethylamino)pyridine, and 38.34 mg (0.2 mMol, 2 equivalents) of N-(3-Dimethylaminopropyl)-N′-ethylcarbodiimide hydrochloride were mixed in 4 ml of dichloromethane, and incubated for 30 minutes at room temperature stirring. 23.3 mg (0.1 mMol, 1 equivalent) of Homopiperazine-benzothiazole amine was dissolved in dichloromethane and added to the aforementioned mixture and allowed to react for 4 hours. The solution was aqueous extracted, dried with anhydrous sodium sulfate, and the product was separated via silica gel flash chromatography with isocratic 5:5 Ethylacetate:dichloromethane. Product was isolated in 90% yield.

(g) **Compound 11**: 15.3 mg (0.1 mMol, 1 equivalent) of 5-cyanothiophene-2-carboxylic acid, 2.44 mg (0.02 mMol, 0.2 equivalents) of 4-(Dimethylamino)pyridine, and 38.34 mg (0.2 mMol, 2 equivalents) of N-(3-Dimethylaminopropyl)-N′-ethylcarbodiimide hydrochloride were mixed in 4 ml of dichloromethane, and incubated for 30 minutes at room temperature stirring. 23.3 mg (0.1 mMol, 1 equivalent) of Homopiperazine-benzothiazole amine was dissolved in dichloromethane and added to the aforementioned mixture and allowed to react for 4 hours. The solution was aqueous extracted, dried with anhydrous sodium sulfate, and the product was separated via silica gel flash chromatography with isocratic 9:1 dichloromethane:Methanol. Product was isolated in 76% yield.

(h) **Compound 12**: 20.7 mg (0.1 mMol, 1 equivalent) 5-Bromo-2-thiophenecarboxylic acid, 2.44 mg (0.02 mMol, 0.2 equivalents) of 4-(Dimethylamino)pyridine, and 38.34 mg (0.2 mMol, 2 equivalents) of N-(3-Dimethylaminopropyl)-N′-ethylcarbodiimide hydrochloride were mixed in dichloromethane, and incubated for 30 minutes at room temperature stirring. 23.3 mg (0.1 mMol, 1 equivalent) of Homopiperazine-benzothiazole amine was dissolved in dichloromethane and added to the aforementioned mixture and allowed to react for 1-4 hours. The solution was aqueous extracted, dried with sodium sulfate, and the product was separated via silica gel flash chromatography with isocratic 9:1 dichloromethane:Methanol as the solvent. Product was isolated in 82% yield.

**Generation of *Dme* HhC Homology Model**

A homology model of *Drosophila melanogaster* HhC was generated from the existing HINT subdomain crystal structure (PDB ID: 1at0) and a generated model of the SRR subdomain, using a SRR model from a published Sonic HhC model [7] as a template.

SRR Model Generation - The template SRR and the *D.me* SRR protein sequences were aligned using ClustalOmega [8]. Subsequently this protein sequence alignment and the 3D Sonic HhC model were used with Modeller 9.25 [9, 10] to create a *D.me* SRR model. The modeled structures that were generated were ranked using MODELLER's internal scoring function, and the model with the lowest (best) score was chosen for full HhC model generation.

Full Model generation and refinement – The published *D.me* HINT domain structure and newly generated SRR structure were aligned to the published full Sonic HhC structure and ligated into a full *D.me* structure using editing tools within Pymol. Using the alignment to the sonic HhC model, cholesterol was placed in a similar binding orientation manually. The full *D.me* structure was submitted to the CHARMM-GUI [11] solution builder input generator, using neutralizing ions, and all other settings default. The output structure was then minimized using GROMACS, with 5000 steps of steepest decent with a Lennard-Jones cutoff radius of 12 Å, followed by 125000 steps of equilibration with all settings default from the CHARMM-GUI GROMACS output.

Small Molecule Docking – The refined *D.me* HhC model and inhibitor structure were prepared for docking by generating pdbqt files using Autodock4 [12]. A grid box encompassing the entire protein model was included in a config file for Autodock Vina[13], choosing an output of 9 docked conformations with all other settings default. All outputs were inspected manually, and the lowest energy docked structure was chosen for the ternary complex model.

### [NMR] Compounds 5 – 12

**Compound 5 – 1H spectra**

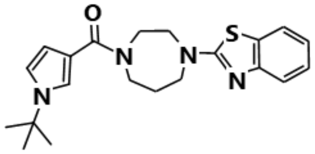

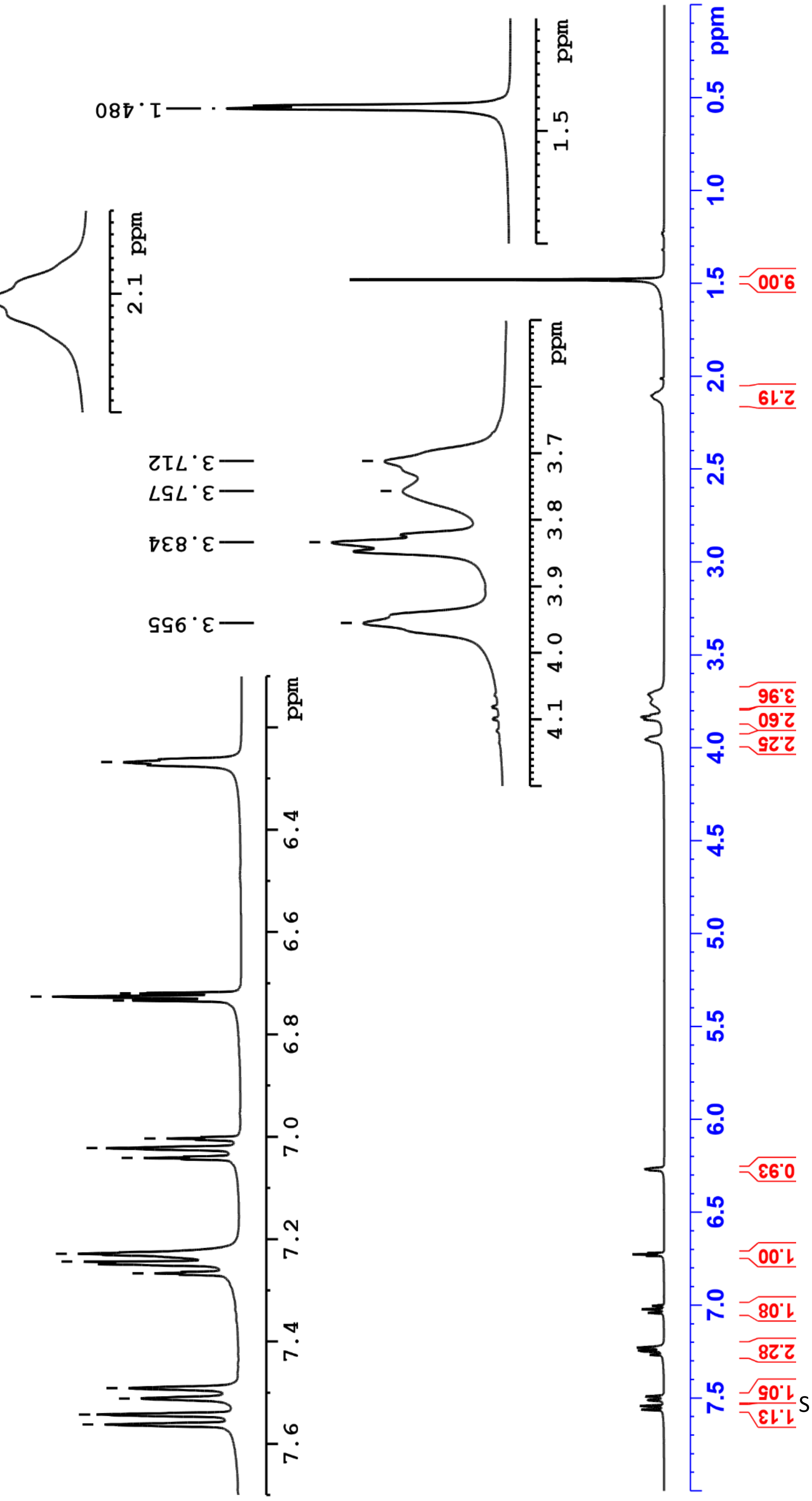

#### Compound 5 – 13C Spectra

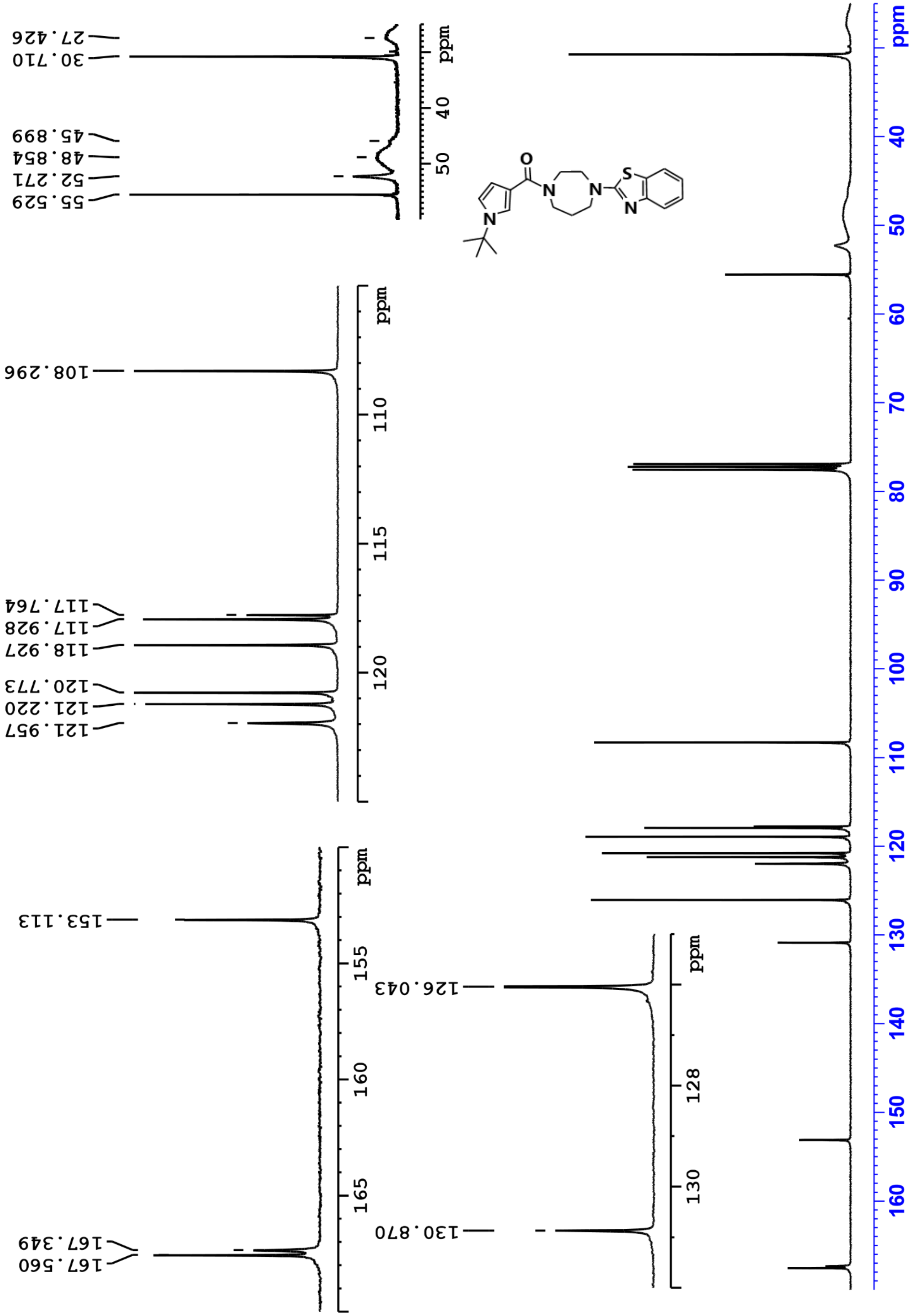

#### Compound 6 – 1H Spectra

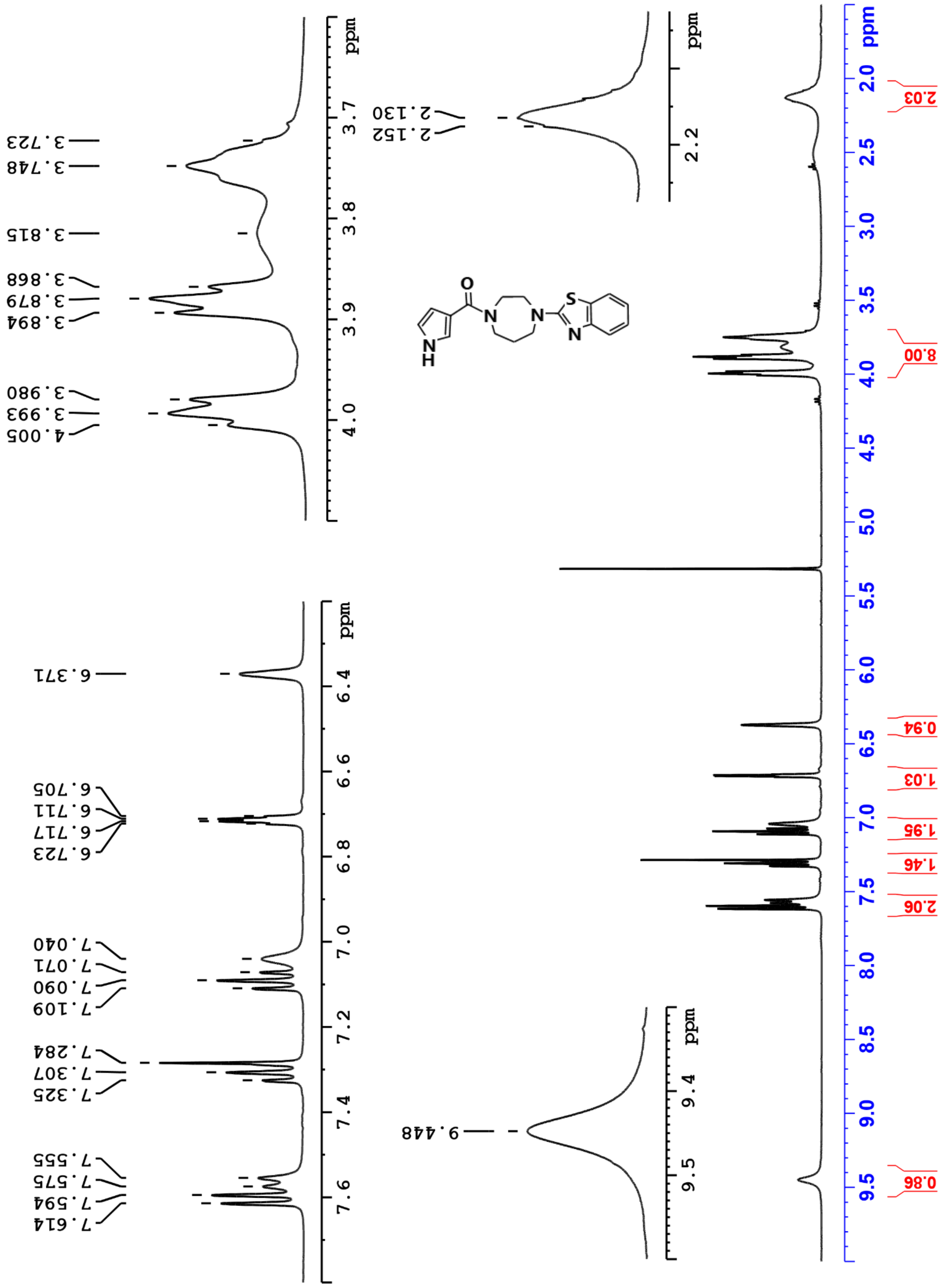

**Compound 6 – 13C Spectra**

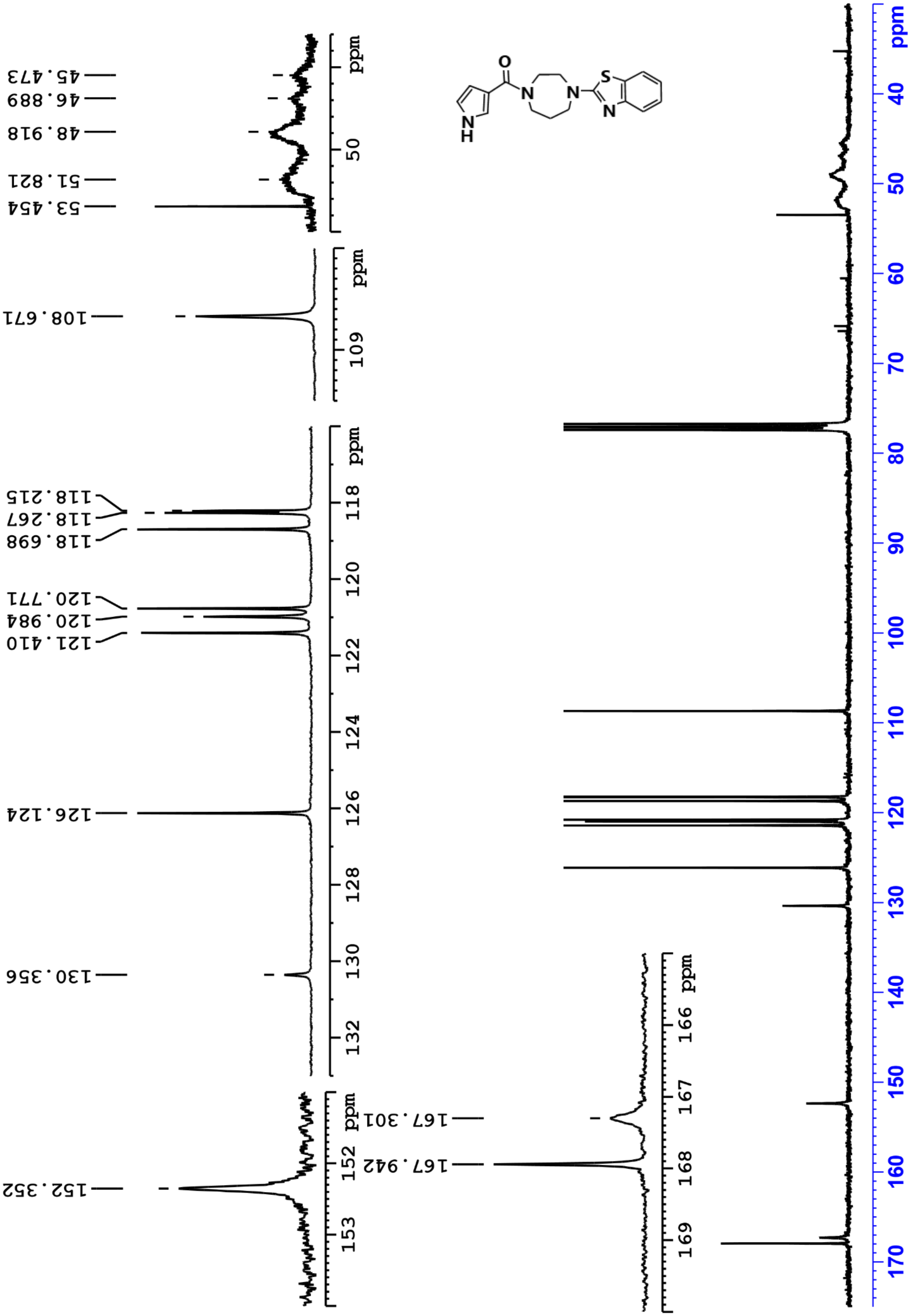

**Compound 7 – 1H Spectra**

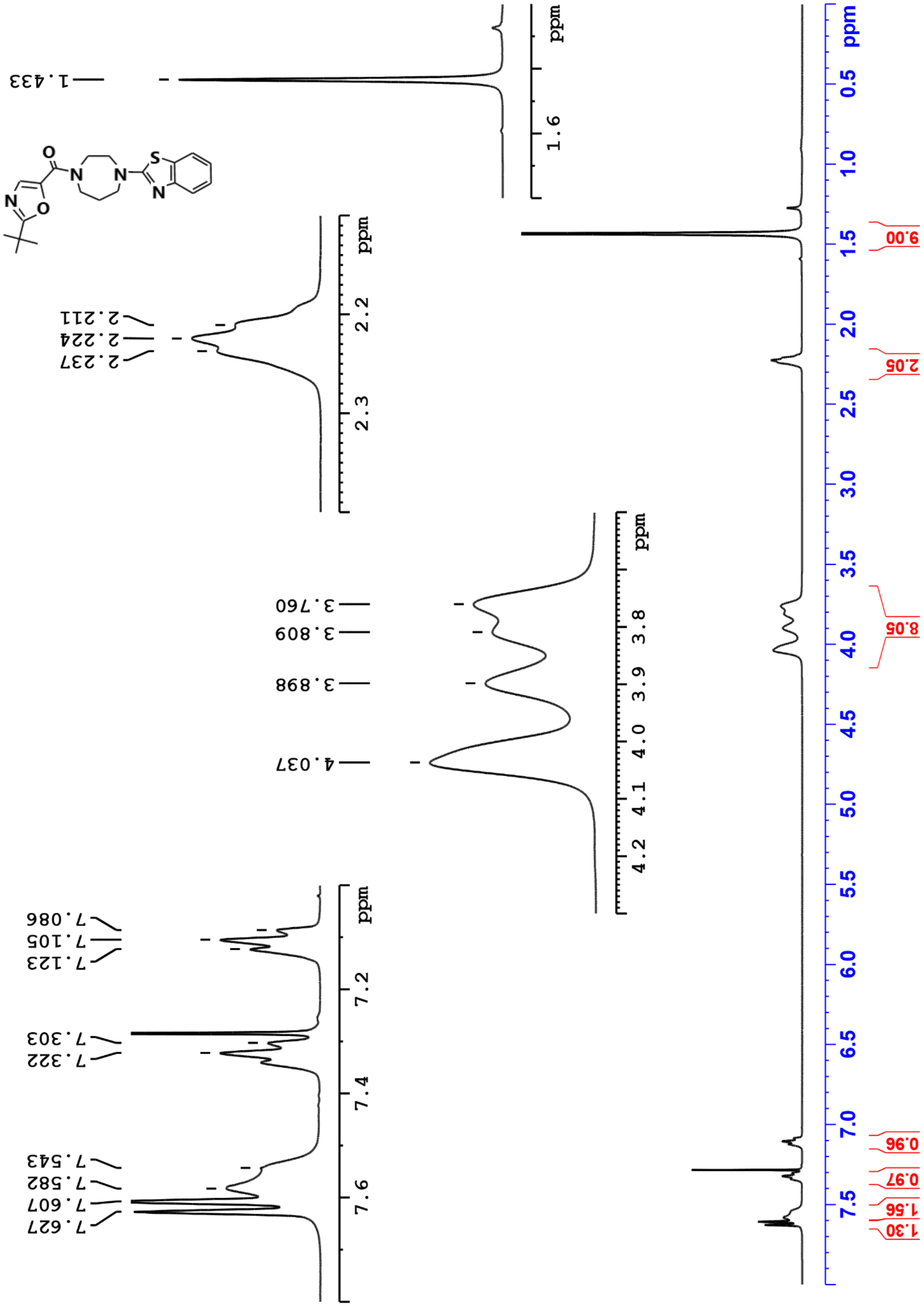

**Compound 7 – 13C Spectra**

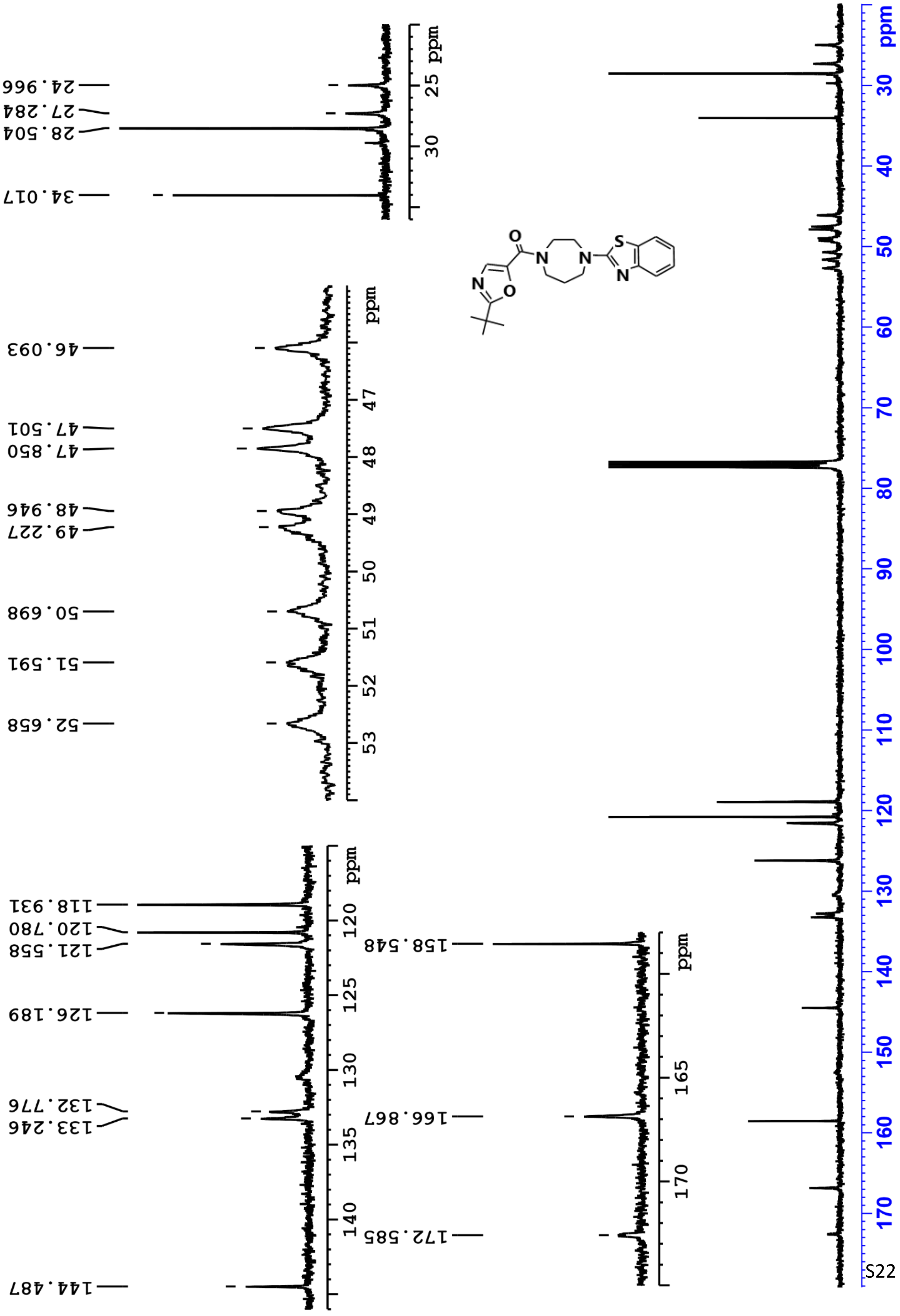

#### Compound 8 – 1H Spectra

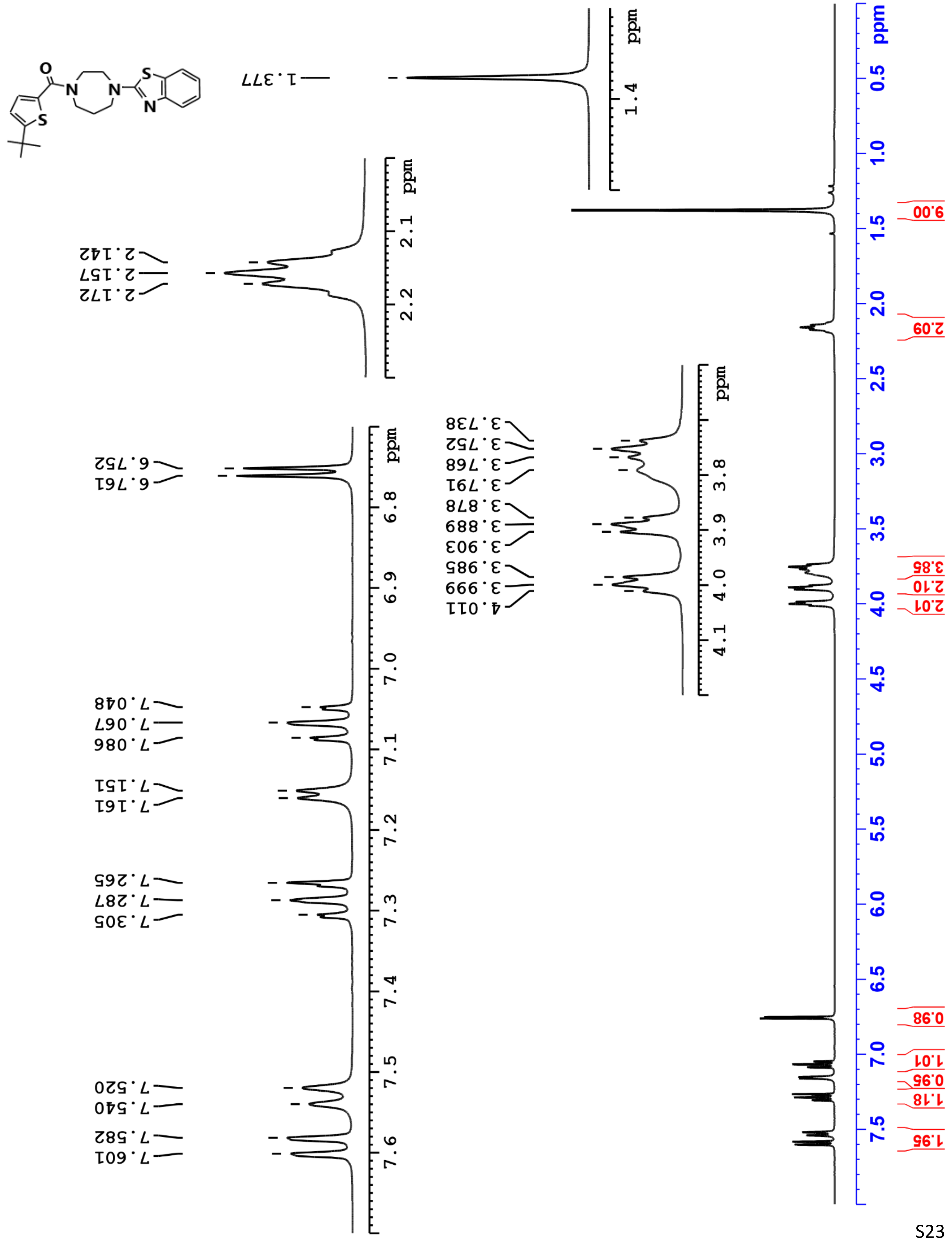

**Compound 8 – 13C Spectra**

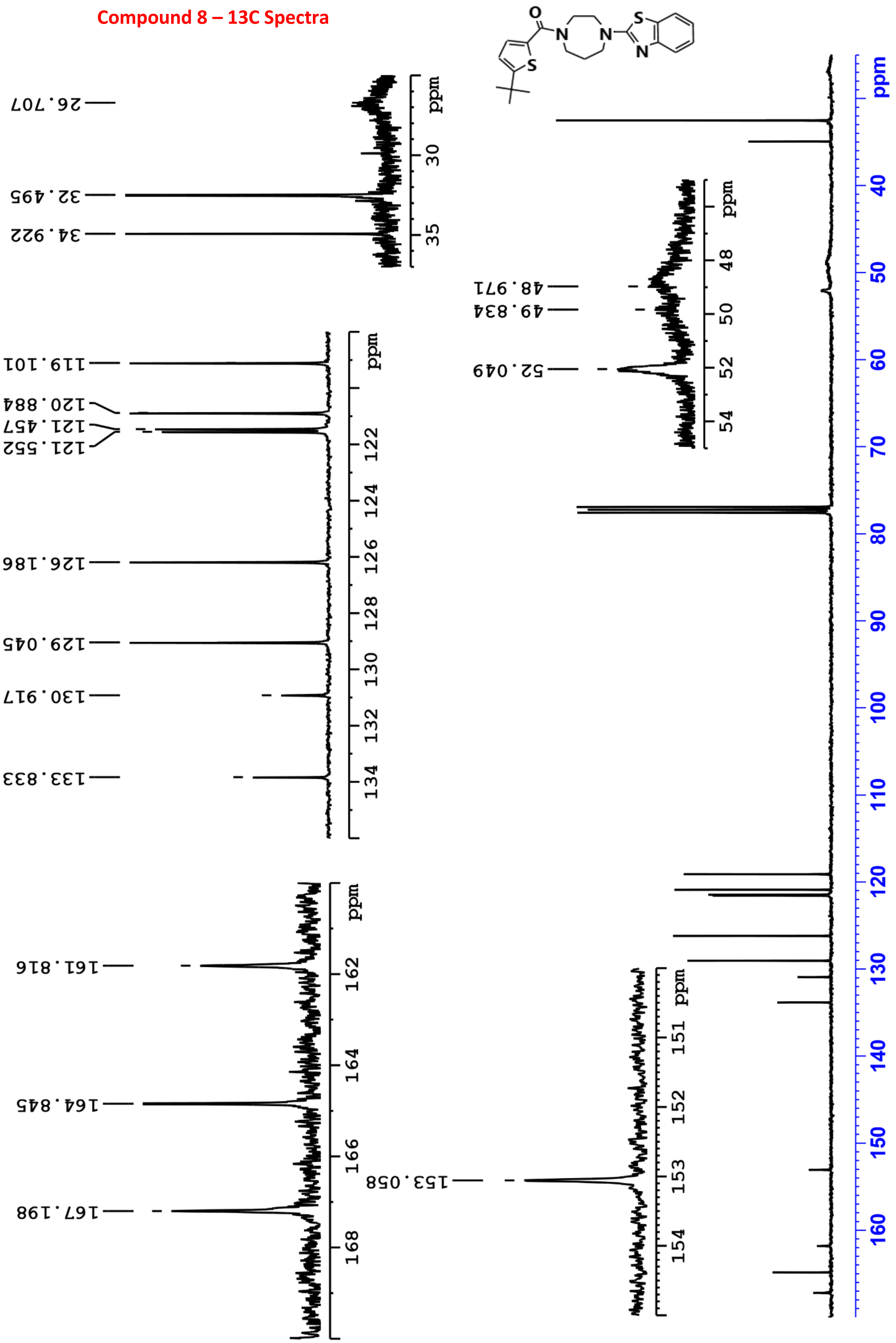

#### Compound 9 – 1H Spectra

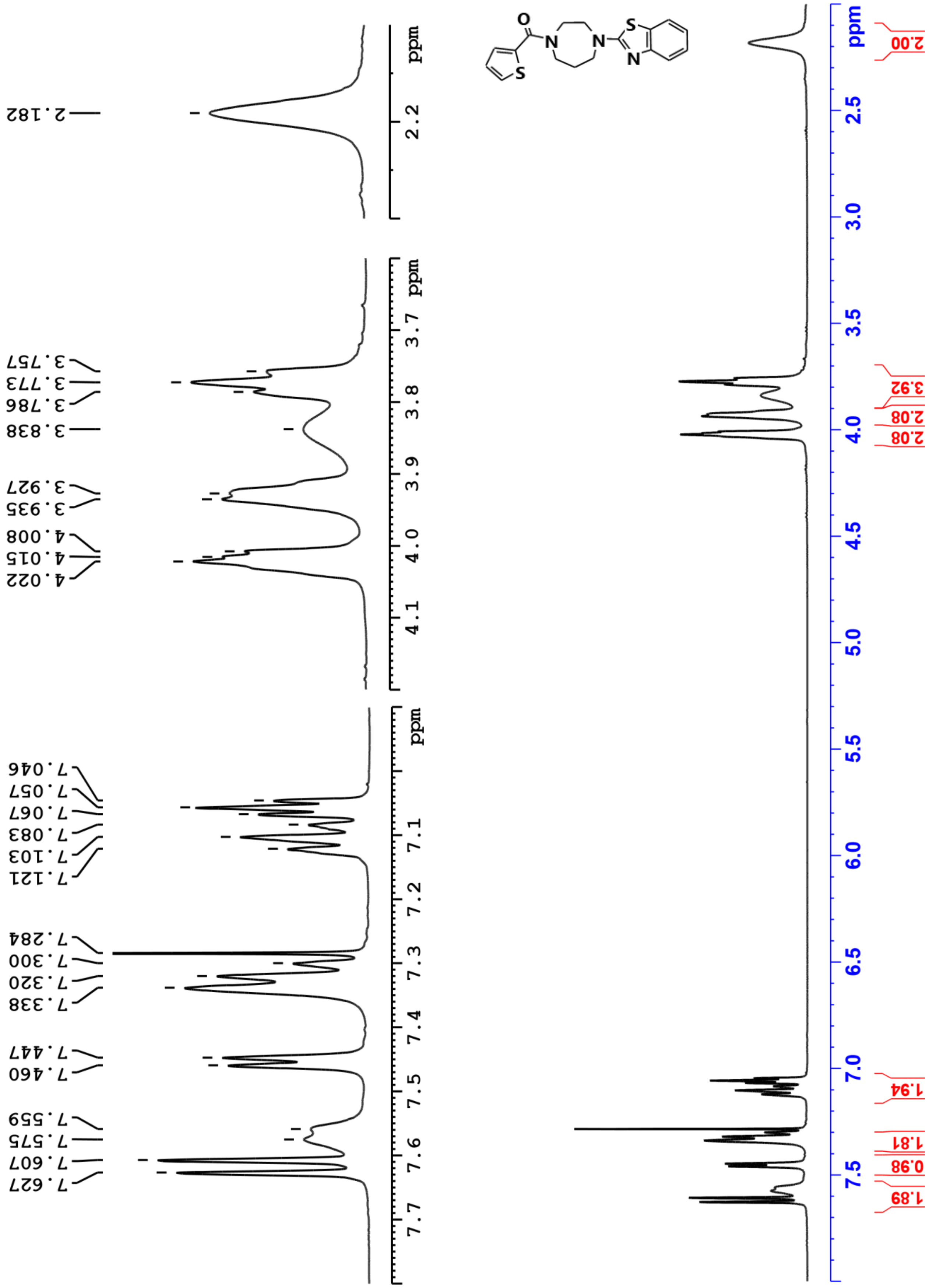

**Compound 9 – 13C Spectra**

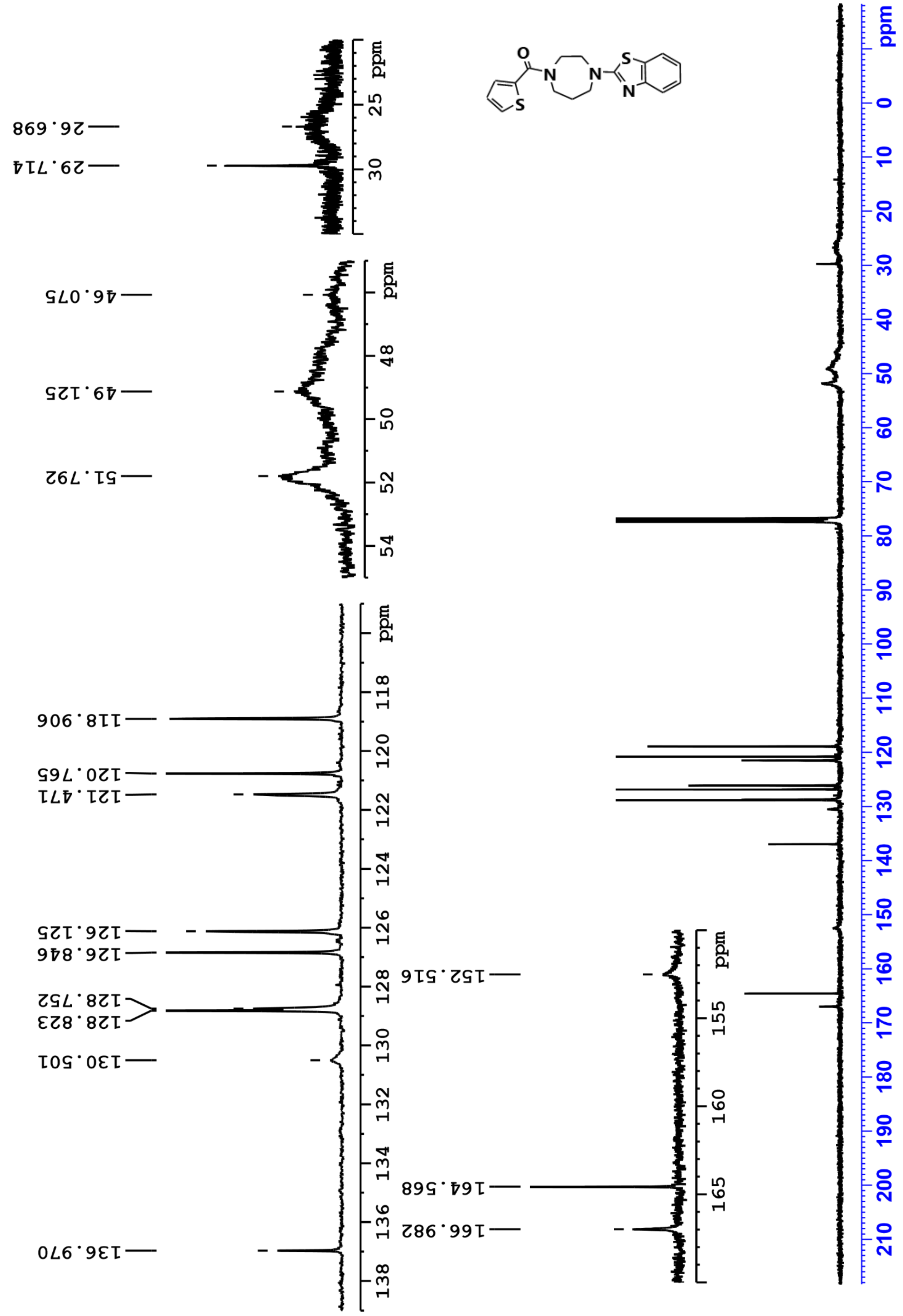

#### Compound 10 – 1H Spectra

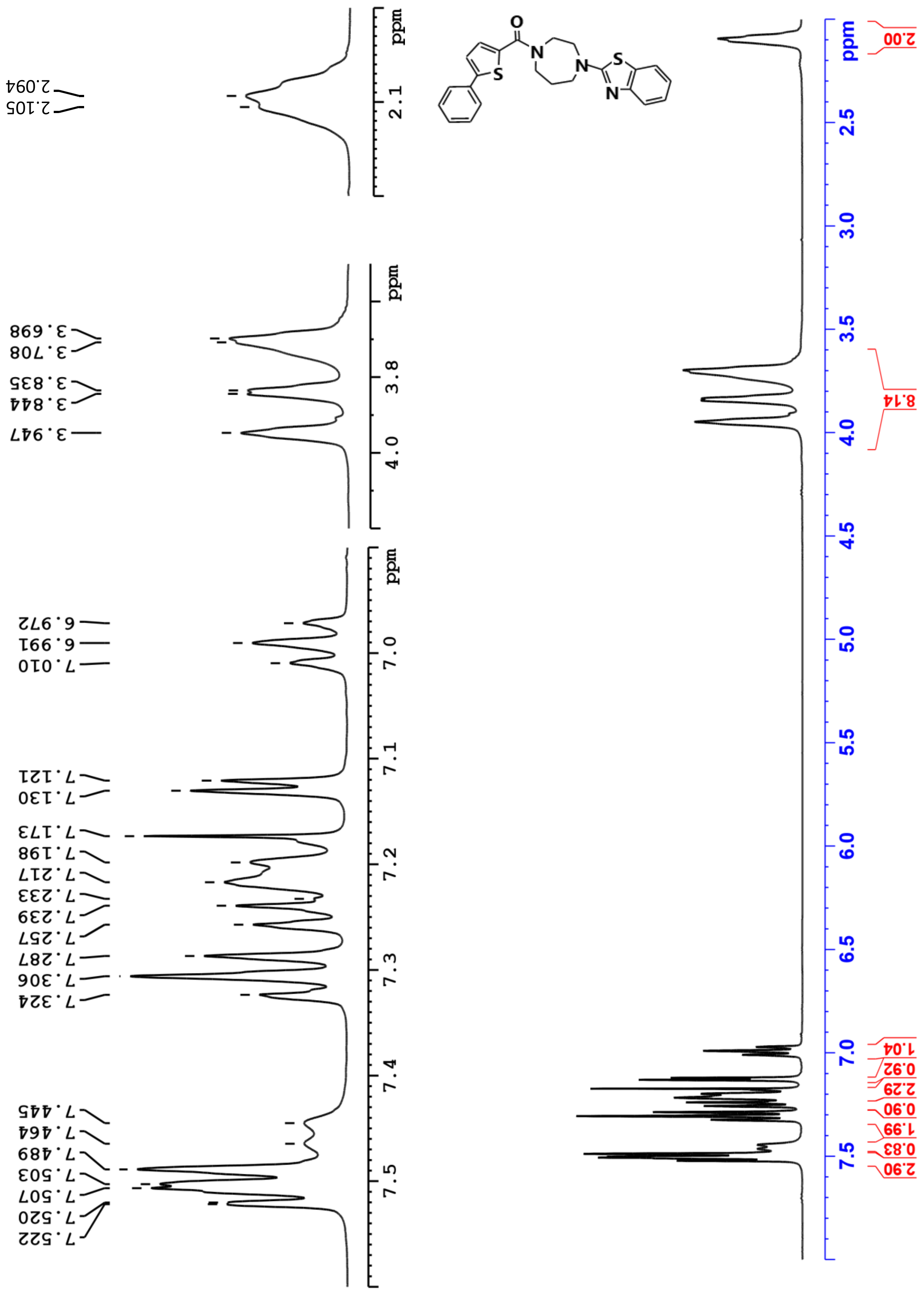

Compound 10 – 13C Spectra

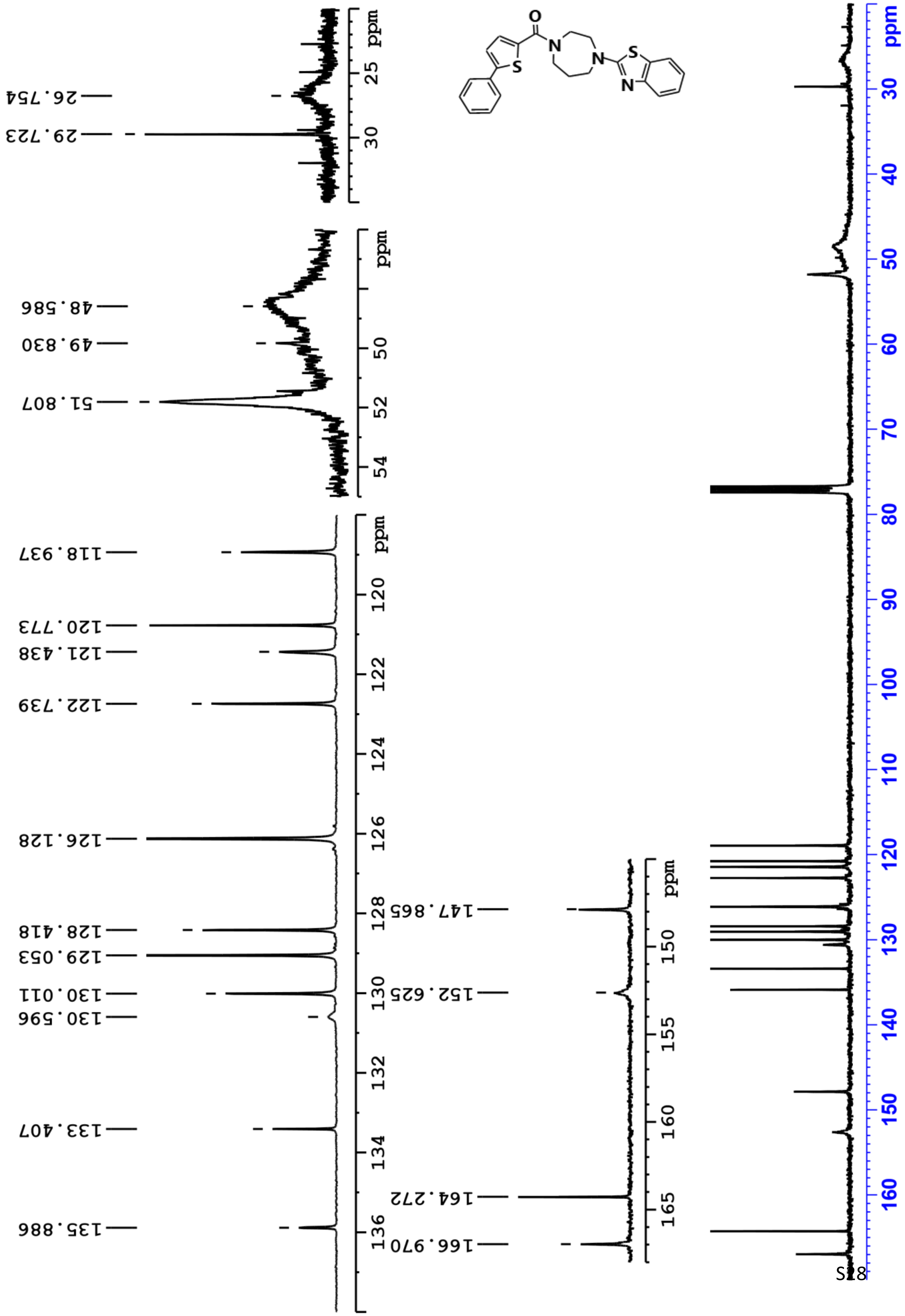

**Compound 11 – 1H Spectra**

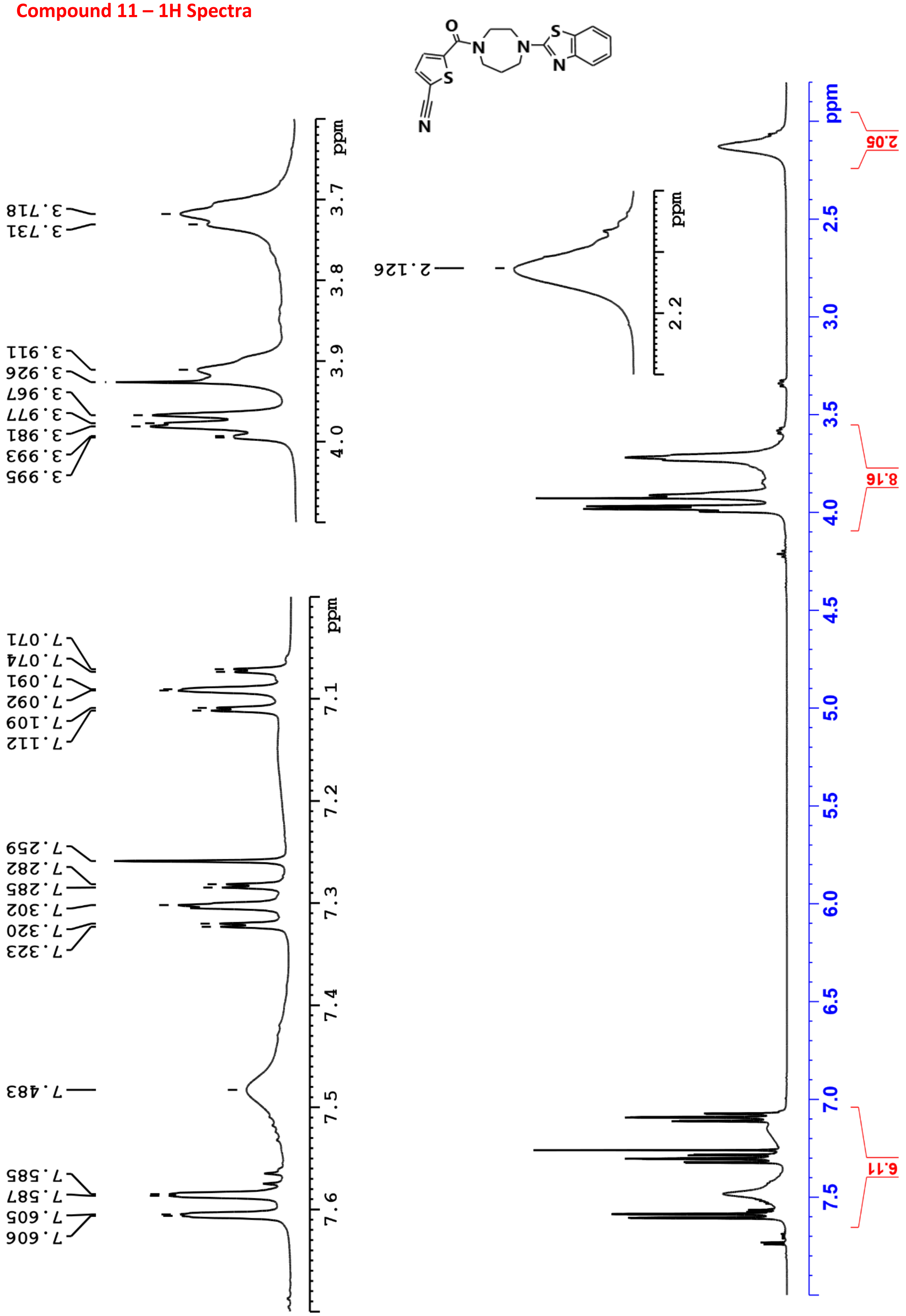

Compound 11 – 13C Spectra

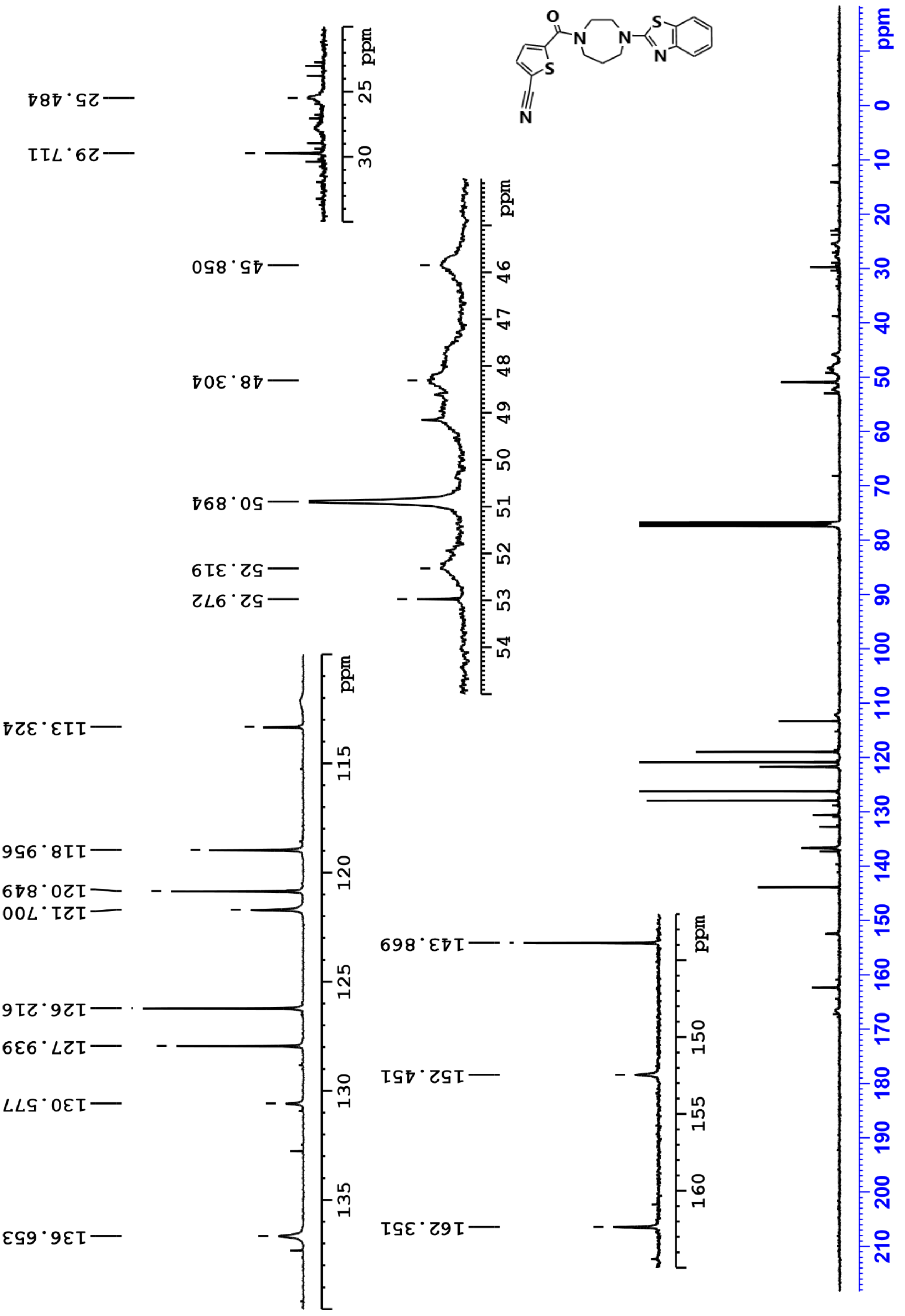

#### Compound 12 – 1H Spectra

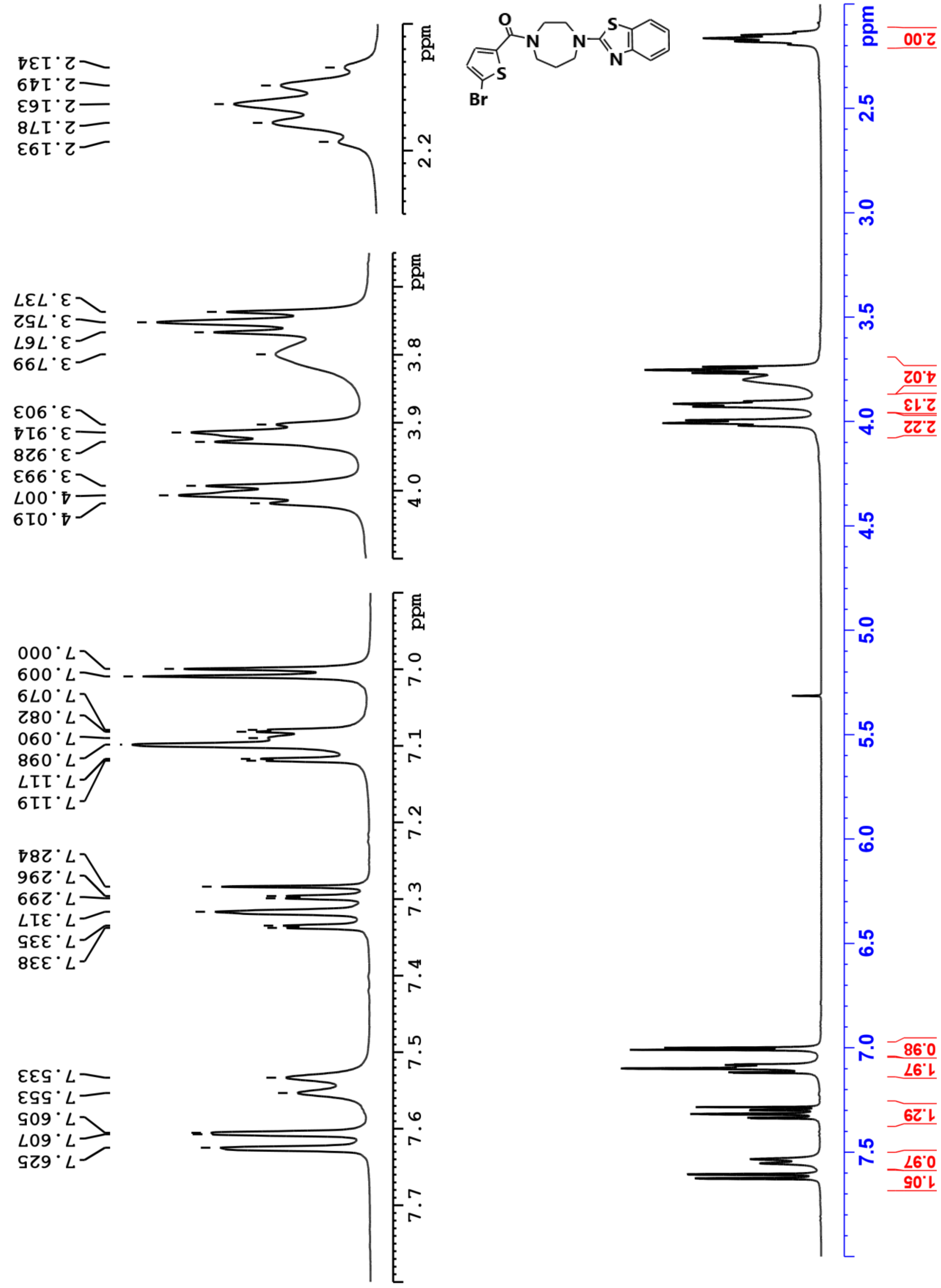

Compound 12 – 13C Spectra

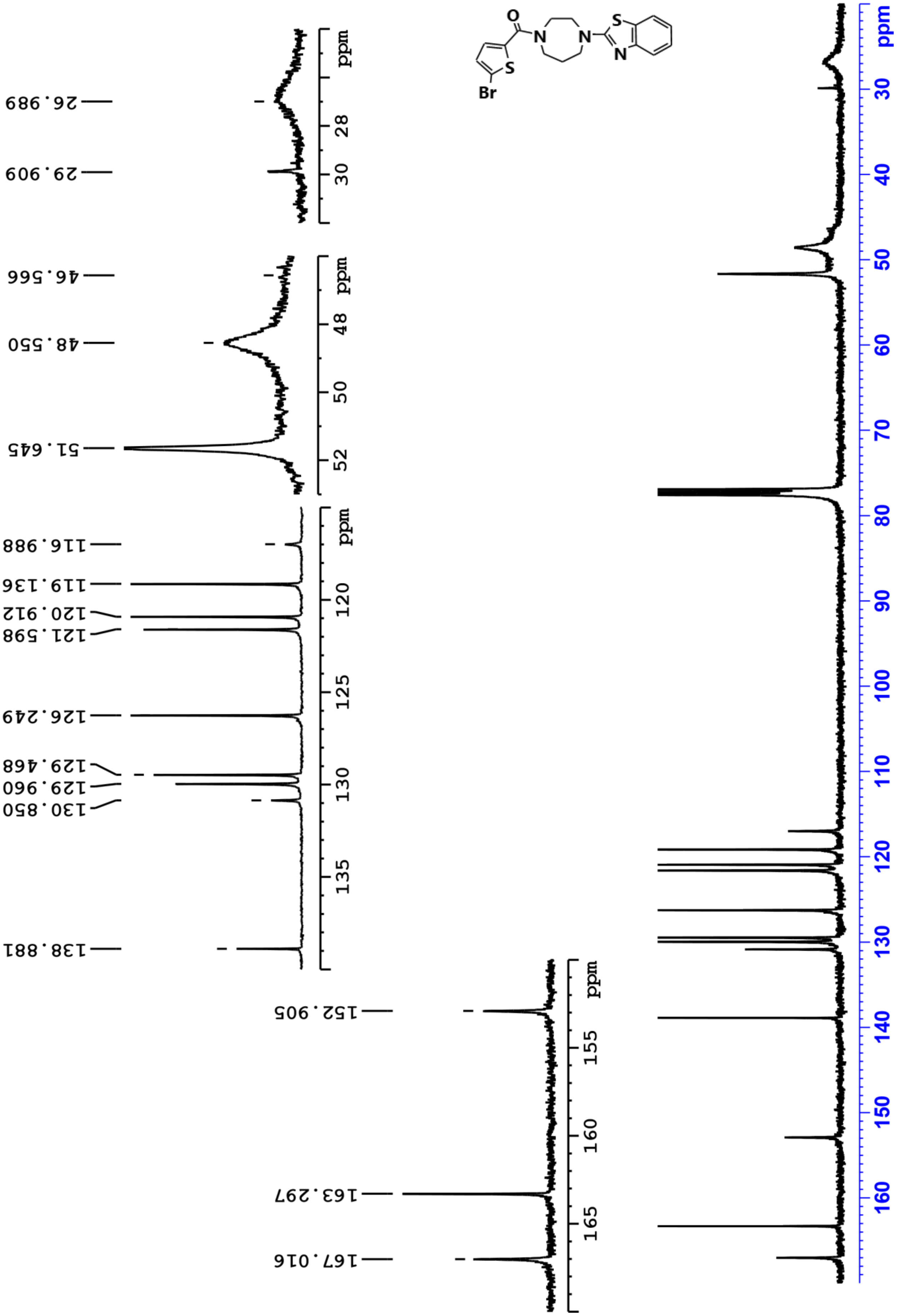

### LCMS

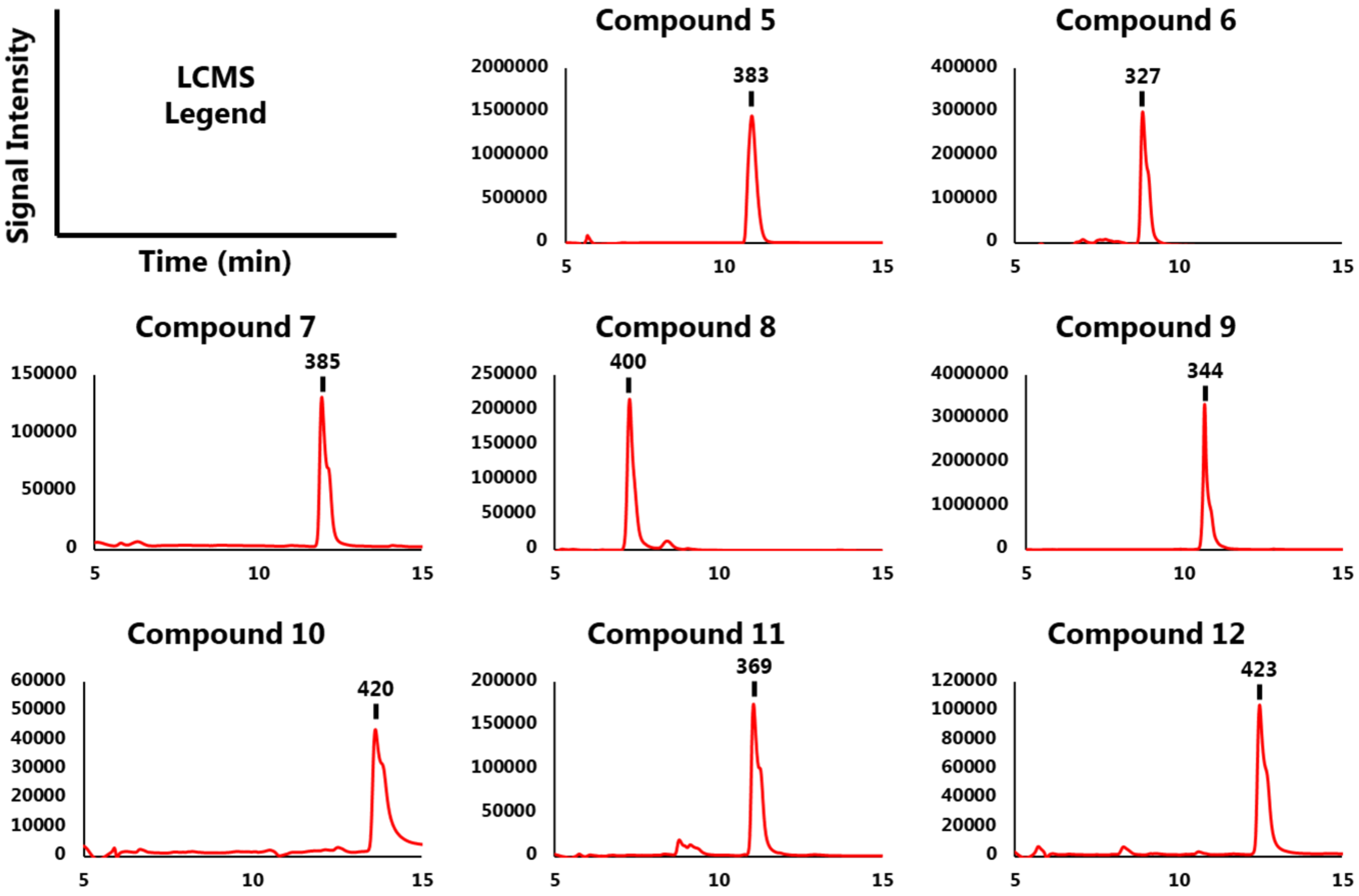


LC traces of synthesized compounds 5-12, with denoted mass of major peaks (g/mol).
